# Robust detection of SARS-CoV-2 exposure in the population using T-cell repertoire profiling

**DOI:** 10.1101/2023.11.08.566227

**Authors:** Elizaveta K. Vlasova, Alexandra I. Nekrasova, Alexander Y Komkov, Mark Izraelson, Ekaterina A. Snigir, Sergey I. Mitrofanov, Vladimir S. Yudin, Valentin V. Makarov, Anton A. Keskinov, Darya Korneeva, Anastasia Pivnyuk, Pavel V Shelyakin, Ilgar Z Mamedov, Denis V Rebrikov, Dmitry M Chudakov, Sergey M. Yudin, Veronika I. Skvortsova, Olga V Britanova, Mikhail A. Shugay

**Affiliations:** Institute of Translational Medicine, Pirogov Russian National Research Medical University, Moscow, Russia; ITMO University, Saint-Petersburg, Russia; Federal State Budgetary Institution “Centre for Strategic Planning and Management of Biomedical Health Risks”, Federal Medical Biological Agency, Moscow, Russia; Department of Genomics of Adaptive Immunity, Shemyakin-Ovchinnikov Institute of Bioorganic Chemistry, Moscow, Russia; Dmitry Rogachev National Medical Research Center of Pediatric Hematology, Oncology and Immunology, Moscow, Russia; Abu Dhabi Stem Cells Center, Abu Dhabi, United Arab Emirates; Skolkovo Institute of Science and Technology, Moscow, Russia; The Federal Medical Biological Agency (FMBA), Moscow, Russia

## Abstract

The COVID-19 pandemic offers a powerful opportunity to develop methods for monitoring the spread of infectious diseases based on their signatures in population immunity. Adaptive immune receptor repertoire sequencing (AIRR-seq) has become the method of choice for identifying T cell receptor (TCR) biomarkers encoding pathogen specificity and immunological memory. AIRR-seq can detect imprints of past and ongoing infections and facilitate the study of individual responses to SARS-CoV-2, as shown in many recent studies. Here, we have applied a machine learning approach to two large AIRR-seq datasets with more than 1,200 high-quality repertoires from healthy and COVID-19-convalescent donors to infer TCR repertoire features that were induced by SARS-CoV-2 exposure. The new batch effect correction method allowed us to use data from different batches together, as well as combine the analysis for data obtained using different protocols. Proper standardization of AIRR-seq batches, access to human leukocyte antigen (HLA) typing, and the use of both α- and β-chain sequences of TCRs resulted in a high-quality biomarker database and a robust and highly accurate classifier for COVID-19 exposure. This classifier is applicable to individual TCR repertoires obtained using different protocols, paving the way to AIRR-seq-based immune status assessment in large cohorts of donors.

## Introduction

Pathogen exposure leaves an imprint on the T cell immunity, with pathogen recognition being guided by antigens presented by HLA molecules and resulting in T cell expansion and formation of immunological memory^1^. This imprint can be explored by sequencing the T cell receptor (TCR) repertoire^2^, and recent studies have revealed that the structure of TCR repertoires derived from peripheral blood mononuclear cells (PBMC) can be explained by a relatively simple V(D)J rearrangement model^3^ together with the signs of past infections^4^. Many studies have utilized adaptive immune receptor repertoire (AIRR)-seq data to infer biomarkers associated with donor phenotype and pathology^5,6^, including efforts directed at predicting exposure to common pathogens such as cytomegalovirus (CMV) or SARS-CoV-2 from repertoire data^7–9^.

The recent COVID-19 pandemic has led to increased effort to analyze TCR repertoires^10,11^. However, most studies have been conducted on small cohorts of individuals (<500 donors), which may not be sufficient for large-scale statistical analysis^12^. Existing studies for large cohorts have shown high-quality classification of COVID-19 severity based on VJ gene segment usage or TCR sequence patterns^13,14^, but the authors did not employ batch effect correction methods, which could affect the generalizability of the results. The challenge in creating a universal pipeline for TCR biomarkers identification lies in tackling prominent batch effects in AIRR-seq data, often characterized by imbalanced V and J gene usage profiles and the existence of common TCR variants caused by cross-sample contamination. Interestingly, traces of the COVID-19 immune response can be identified in both healthy donors and patients^15^, which may indicate cross-reactivity of the pre-existing memory T cell immune response to SARS-CoV-2. In addition, the virus is rapidly evolving^16–18^, and it has recently been shown that the robustness of the SARS-CoV-2-specific immune response is mediated by the diversity of the antigen-specific T cells^19^. Notably, the immune response is also strongly differentiated by the patient’s HLA alleles^5^, which is important to account for when building AIRR-based classifiers.

In the present study, we report a classifier that can distinguish SARS-CoV-2-exposed patients from uninfected donors, including individuals sampled before the COVID-19 pandemic. The main idea of this approach is to use complementarity-determining region 3 (CDR3) sequences, which are highly diverse and encode most information on TCR antigen specificity, as biomarkers for training the classifier, extending far beyond the set of previously reported TCRs recognizing cognate SARS-CoV-2 epitopes^8,20,21^.

Our cohort features individuals of known COVID-19 status and HLA class I and II haplotype that were subject to deep profiling of both TCRα and β chain repertoires. We extracted CDR3 sequences that can serve as biomarkers of COVID-19 status and assessed them by studying HLA association and TCRα-β chain pairing, and discovered a large number of homologous groups of TCR sequences (‘metaclonotypes’) similar to known SARS-CoV-2-related TCRs. We aimed to minimize batch effects that are prominent in repertoire-sequencing studies, thereby allowing us to generalize our classifier across data from different batches and remove potential biases^22–24^, resulting in a robust classifier that shows good performance on unseen data batches. We also used a previously reported cohort of convalescent and healthy individuals from Adaptive Biotechnologies^8,25^ to further validate the performance of our classifier.

## Results

### Study design overview

The main aim of our study was to find TCR sequence biomarkers and develop a bioinformatic pipeline that allows the construction of an accurate and robust classifier that distinguishes COVID-19-convalescent donors from unexposed individuals, as summarized in **Figure 1**. We performed immunosequencing of the rearranged TCRα and β regions from donor PBMCs. For the cohort described in this study (Cohort I), we sequenced both chains of the TCR heterodimer, as both chains are required to properly predict antigen recognition^26^. We ran a conventional T cell repertoire data analysis pipeline (see **Methods**) and pre-processed the data to remove low-coverage samples.

**Figure 1.**
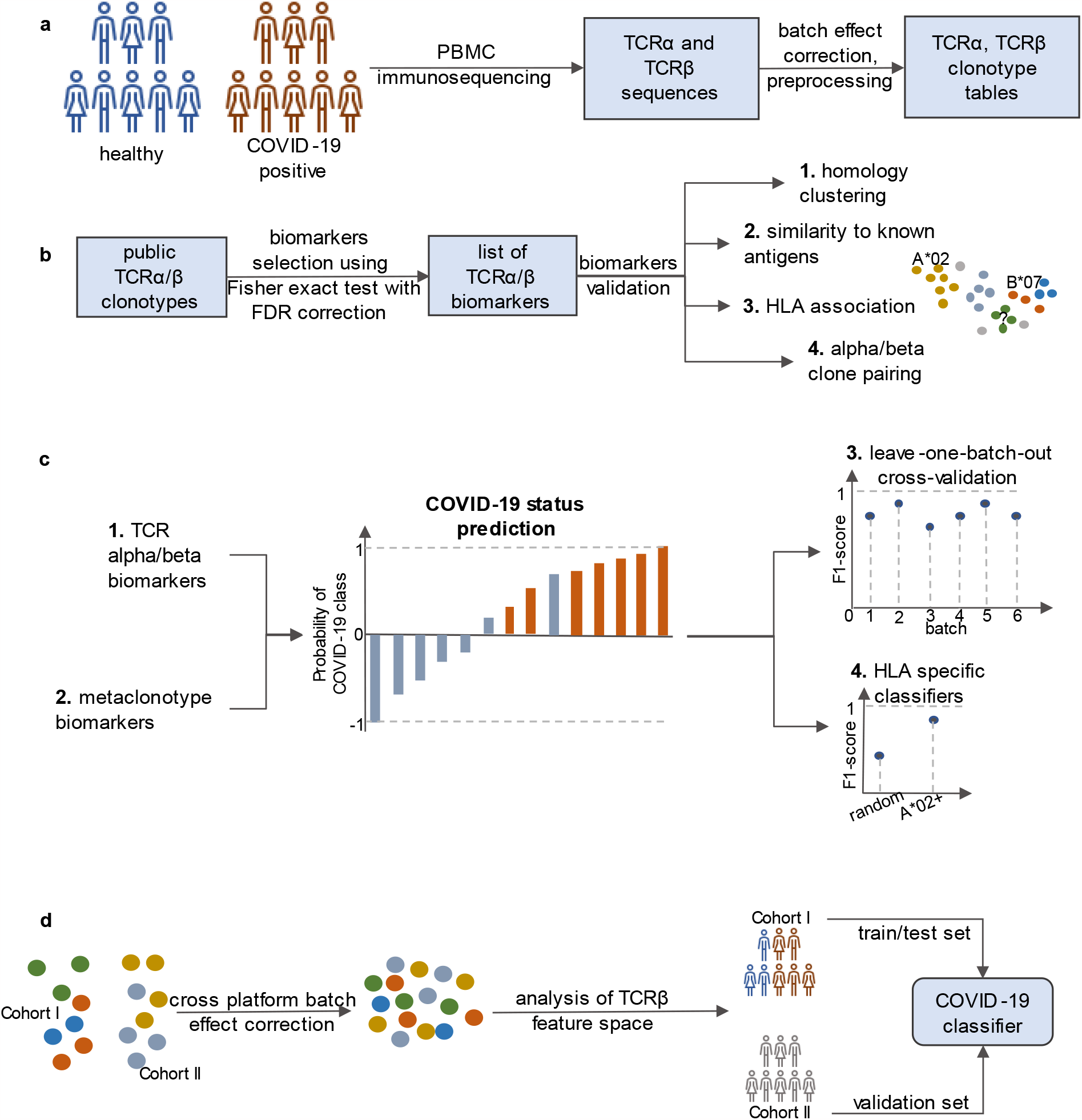
Experimental and analytical procedure. **A**. Peripheral blood samples were obtained for donors from Cohort I with known COVID-19 status and HLA context. TCRα and β loci were sequenced using a conventional protocol involving multiplex PCR (see **Methods**). Samples were preprocessed and mapped to identify V/D/J alleles and extract CDR3 regions. A batch-effect correction procedure was applied to normalize V(D)J rearrangement frequencies. **B**. TCRα and β biomarkers were selected for both T cell chains separately using the Fisher exact test. These biomarkers were aggregated by clustering TCR sequences into ‘metaclonotypes’. Annotation of these clusters was performed using a database of TCR sequences with known antigen specificity, association with HLA metadata, and the results of α-β chain pairing analysis. **C**. Classifiers including various sets of features were constructed based on metaclonotype biomarkers described above. These classifiers were trained and evaluated using a leave-one-batch-out cross-validation technique. **D**. We used another previously published cohort (Cohort II)^8, 25^ to assess the robustness of our classifier. Batch-effect correction and sample preprocessing techniques were applied to both cohorts, and the classifier was trained entirely on one of the cohorts and validated on the other.

After sequencing, we processed the data to extract clonotype tables from each sample — that is, TCR sequences rearranged from specific V and J genes and having a specific CDR3 region sequence. Hereafter, we will focus on CDR3 amino acid sequences as a proxy for specific T cell clonotypes, and refer to the number of sequencing reads corresponding to that clonotype as the clonotype size. Of the samples from Cohort I which passed the read-count threshold, 383/377 TCR α/β samples were from healthy donors (SARS-CoV-2 PCR test negative or obtained prior to pandemic) and 890/848 were from COVID-19-positive patients (**Supplementary Figure 1A, B**). The majority of samples were accompanied by information on HLA class I and II alleles. Samples were prepared and sequenced in nine batches.

We proceeded to select a set of CDR3 sequences that can serve as biomarkers and form a feature list for COVID-19 status classifier. Selecting an unbiased set is problematic due to the batch effects that stem from cross-sample contamination^27^ and method-specific biases^23^. Such biases can also affect the classifier training procedure, as it can learn from them when batches are unbalanced in terms of cases. In order to tackle such batch effects, we performed a standardization procedure that allowed us to correct for differences in V and J gene usage, which are the major source of differences across batches. We also validated the resulting set of clonotypes in several ways. Co-occurrence of specific TCRα and β clonotypes can serve as an independent validation for biomarkers and their co-association with some specific pathogen. Additional information on donor HLAs was also provided to filter the set of biomarkers based on HLA restriction; association with donor HLA serves as additional evidence for TCR specificity to a specific set of antigens presented in a given donor, and allows detection of the fingerprints of past and present infection^28^. Furthermore, clonotypes with similar sequences can be aggregated into ‘metaclonotype’ biomarkers based on clonotype graph analysis. Finally, we trained various COVID-19 status classifiers on selected batches derived from Cohort I data using different algorithms and incorporating different feature sets. We verified the robustness of our results using independent batches from Cohort I as well as data from the previously published Cohort II that contain TCRβ sequences from COVID-19-convalescent donors^8^ merged with control samples obtained prior to the pandemic^25^ (**Supplementary Figure 1C, D**).

### Batch effects and their correction in TCR repertoires

As with many large-scale studies involving hundreds of subjects, our dataset was prepared in several batches. These batches were uneven both in terms of the number of subjects and imbalances in the ratio of COVID-19-convalescent patients to healthy donors (**Figure 2A; Supplementary Figure 2A, B**). Since TCR repertoire sequencing data is extremely high-dimensional and complex, even minor differences in library preparation and sequencing between batches can lead to biases in the comparative analysis of repertoires and obfuscate COVID-19-related features. We therefore utilized V and J gene frequency profiles (*i*.*e*., ‘gene usage’) of repertoires to monitor for these biases and biological effects.

**Figure 2.**
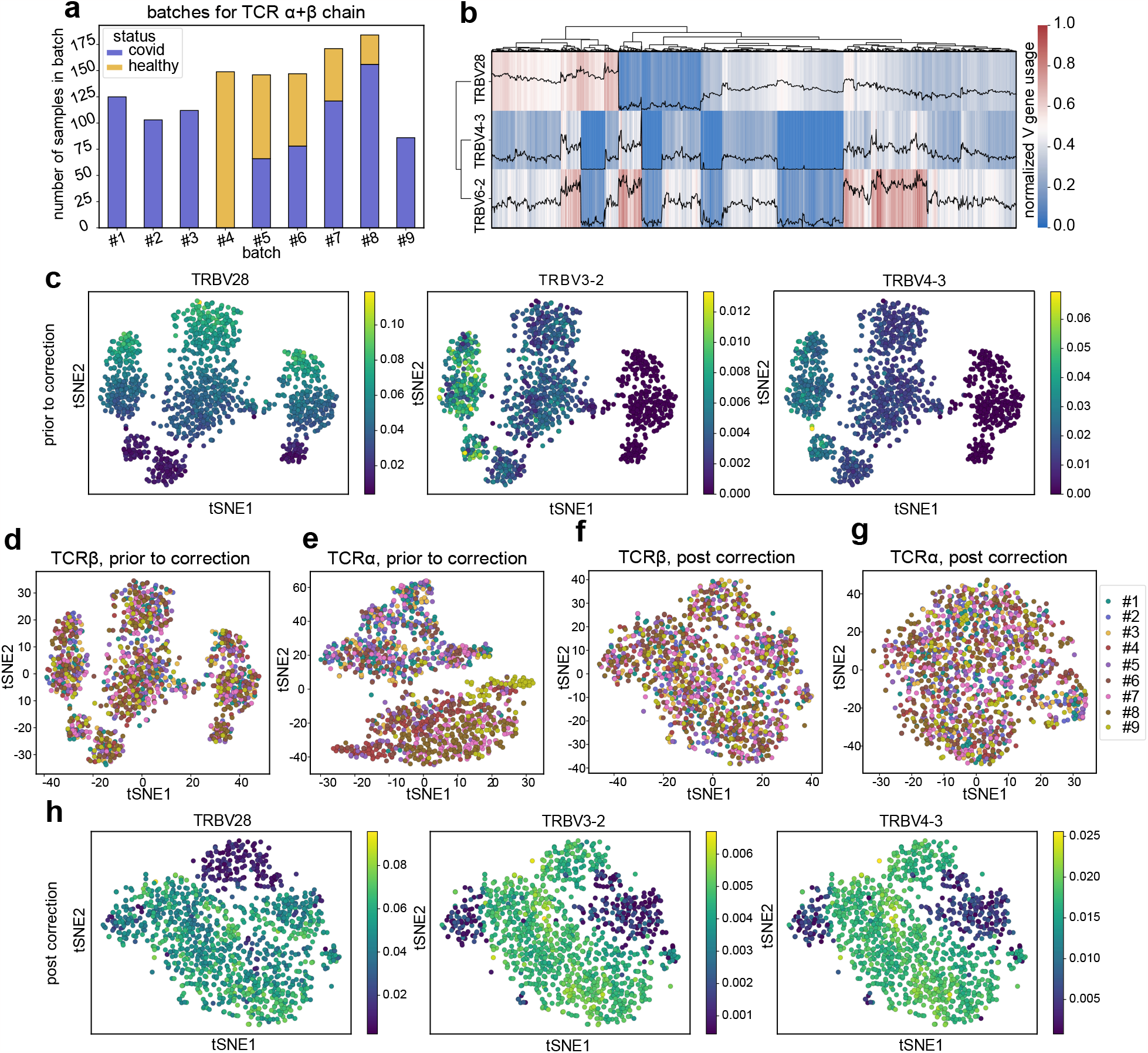
Batch-effect and genotype-related differences in V gene usage. **A**. A bar plot of total number of subjects and the number of COVID-19-convalescent and healthy cases for nine donor sample batches from Cohort I. Numbers reflect samples that passed the sequencing depth cutoff for both TCRα and β chains. **B**. Heatmap of gene usage, showing the imprint of TRBV28/3-2/4-3 haplotypes in the dataset. Black lines show gene usage Z-scores for each of the three genes. **C**. t-SNE plots of TRBV usage according to the frequency of TRBV28, TRBV3-2 and TRBV4-3 gene usage. **D, E**. Visualization of batch effects using t-SNE. Plots show similarity of TRBV (**D**) and TRAV (**E**) gene usage profiles between samples. Samples are colored by batch. **F, G**. Same as **D, E**, but for TRBV (**F**) and TRAV (**G**) gene usage profiles after batch-effect correction. **H**. t-SNE plots of TRBV usage for the batch effect corrected data according to the frequency of TRBV28, TRBV3-2 and TRBV4-3 gene usage.

We start with V gene usage clustering analysis, and used standard multidimensional reduction approaches such as principle component analysis (PCA) and t-stochastic neighbor embedding (t-SNE) for visualization. From the gene usage heatmap analysis, one can spot a prominent and peculiar pattern of TRBV28, TRBV6-2, and TRBV4-3 co-usage (**Figure 2B**). This pattern can be attributed to germline differences in the TCRβ locus^29^: there is a well-known 21-kb deletion polymorphism involving both TRBV6-2 and TRBV4-3, while TRBV28 is involved in a single-gene deletion polymorphism. As can be seen in the t-SNE plot from **Figure 2C, D**, there is a three-cluster pattern in TRBV usage that seems to be unrelated to differences between batches. Further analysis showed that this is related to TRBV28 deletion in component 1 and TRBV6-2/TRBV4-3 deletion in component 2 (**Figure 2C**). Ideally, when correcting for batch effects, one should still be able to recover biological differences between samples such as the aforementioned germline polymorphisms. On the other hand, TRAV usage demonstrated clear batch-related clustering, with samples from batches 1 and 8 predominantly located on the positive and negative sides of component 1 in the t-SNE plot (**Figure 2E**). Batch effects were also detected for TRAJ and TRBJ genes (**Supplementary Figure 3**); for both chains, batches sequenced later in time were clustered together (*e*.*g*., batches #8 and #9; **Supplementary Figure 2C, D**). Allelic effects were also visible for the TRBJ, TRAV and TRAJ genes, but we didn’t find any previous reports for these (**Supplementary Figure 3**), which we attribute to the fact that most AIRR-seq studies have focused on the β chain.

We proceeded to perform the batch-effect correction procedure described in detail in the **Methods** section. Briefly, we aimed to normalize gene usage profiles across samples by using Z-scores calculated across each batch for log-transformed gene frequencies. These values were then transformed and re-scaled to reflect the global gene usage hierarchy of the entire dataset. As one can see from **Figure 2F** and **G**, and **Supplementary Figure 2E** and **F**, such normalization negates batch effects such that the samples are uniformly distributed on the t-SNE plot. On the other hand, our correction procedure leaves biological variability intact, as can be seen for the case of TRBV28, TRBV6-2, and TRBV4-3 shown in **Figure 2H**. The biological variability is also preserved for TRAV and TRAJ genes as well as for TRBJ gene usage (**Supplementary Figure 3**). After correcting gene usage profiles, one can re-sample the clonotypes in the repertoires as described in **Methods** and proceed with comparative analysis of donor repertoires to search for COVID-19-associated clonotypes. We trained COVID-19 status classifiers based on V-J gene usage profiles alone, but did not succeed (**Supplementary Figure 4**).

### Identification of COVID-19-associated TCR α and β clonotypes

We proceeded to perform statistical inference of COVID-19-associated TCR clonotypes from batch-corrected repertoires. Inspired by previous work identifying CMV-associated TCR sequences^25^, we performed the Fisher exact test for the contingency table of presence/absence of a given TCR and positive/negative COVID-19 status with one critical modification described below. Note that at this step we considered all batches in our analysis: due to extremely high diversity of TCR repertoires and sparse overlap, each batch will likely bring many novel biomarkers.

First, we made a short-list of candidate TCR α and β clonotypes that were rearranged at least twice—*i*.*e*., supported by at least two unique nucleotide variants or found in at least two samples in our entire dataset. After computing P-values for the association, we ran false-discovery rate (FDR) correction for multiple testing to select significant hits at q = 0.05. Surprisingly, no significant hits were found for TCRβ clonotypes. We suggest that this is due to limited sampling depth of individual repertoires and low cell counts of individual TCR variants in a donor. We then modified our method to allow a single mismatch in the CDR3 amino acid sequence when searching for a clonotype of interest in the repertoire. This adjustment is justified by the fact that a single mismatch in CDR3 doesn’t change antigen specificity most of the time^30^. The other rationale is that the estimate of clonotype population frequency computed this way is in perfect agreement with theoretical baseline V(D)J rearrangement probability^3^. After this adjustment, many significant hits were obtained at q = 0.05. For the final list of COVID-19-associated TCRs, we used the q = 0.01 threshold, arriving at 4,887 and 574 significant TCRα and β biomarkers, respectively (**Supplementary Figure 5A** and **B**). This nearly 10-fold difference in the number of variants for TCRα and β can be explained by the fact that TCRβ clonotypes are less public and more dissimilar due to the presence of D gene segments; as such, finding a significant association for them is harder and requires more samples than for TCRα.

Basic analysis of the incidence rate of COVID-19-associated clonotypes showed that they constitute a detectable fraction in COVID-19-convalescent donor samples (24.1% and 6.8% on average of all clonotypes in TCRα and TCRβ repertoires respectively; **Supplementary Figure 5A, B, C, D**) that is significantly higher compared to healthy donors (P = 1.2 x 10^-33^ and 2.6 x 10^-15^ for TCRα and β respectively, Mann-Whitney test; **Supplementary Figure 5E, F)**. Manual inspection of clonotype lists revealed that some of the found clonotypes had spurious CDR3 sequences that are either extremely short or too long, or non-canonical (*e*.*g*., CDR3 not starting with Cys and ending with Phe/Trp). We used the V(D)J rearrangement model^31^ to compute the generation probability for each of the clonotypes in our list.

We found outliers both for TCRα and β sequences (**Supplementary Figure 5G, H**) that match manually-detected spurious sequences and removed them, leaving 4,393 and 567 TCRα and β biomarkers, respectively, for further analysis. The results remained unchanged after spurious clonotype removal (**Figure 3A, B, C** and **D**). In order to check the robustness of our results to batch removal, we repeated the analysis while leaving out a validation batch (batch #6), and we were nevertheless able to reproduce the results above. There was a detectable number of rearrangements matching COVID-19-associated TCRs in convalescent donors (**Supplementary Figure 6A, B**), and those were significantly more frequent than in healthy donors (P < 10^-4^ for both α and β, Mann-Whitney test, **Supplementary Figure 6C, D**).

**Figure 3.**
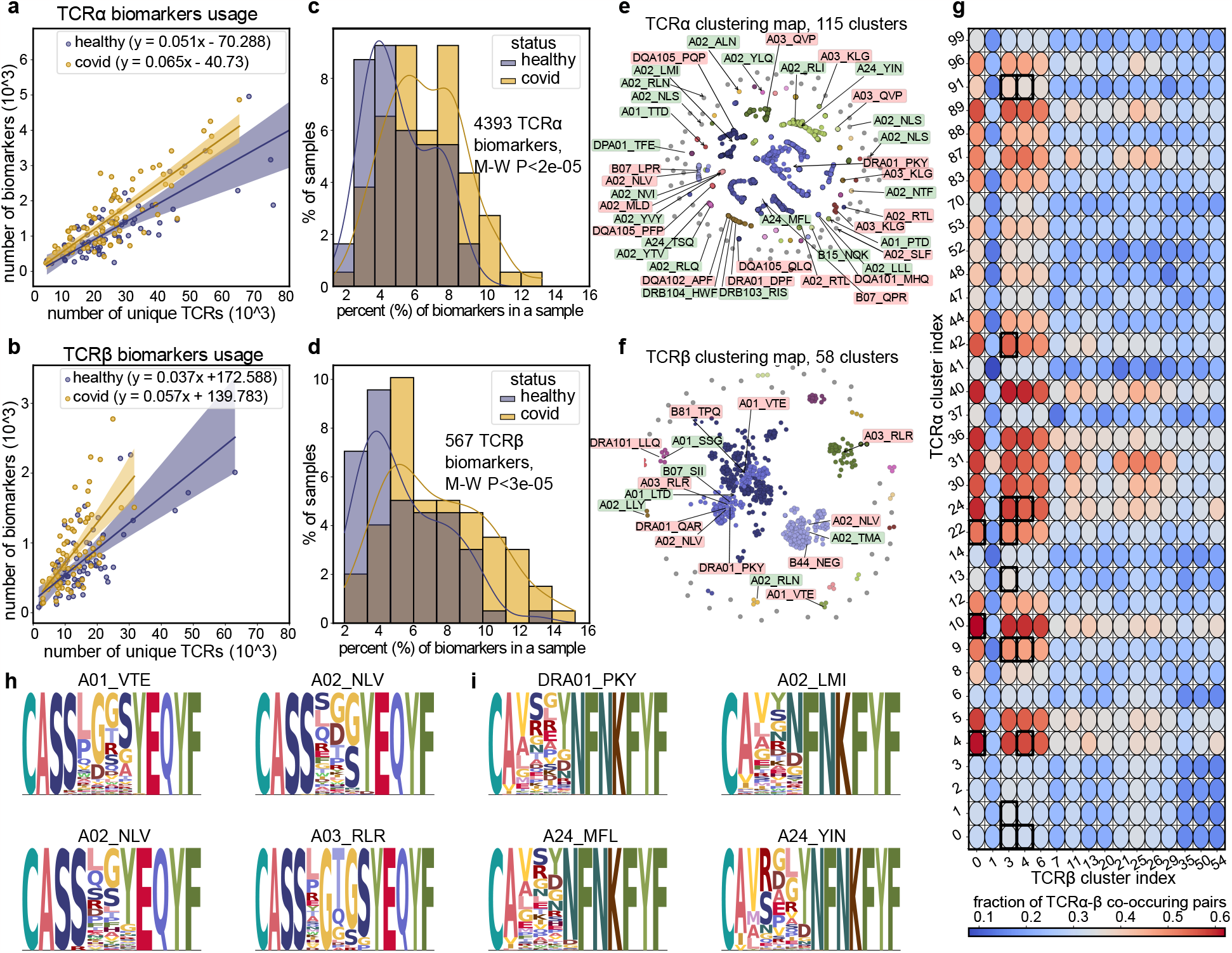
Assessment of COVID-19 TCR α and β biomarkers. **A, B**. Scatter plot showing the number of COVID-19-associated TCRα (**A**) and TCRβ (**B**) clonotypes (Y axis) plotted against the total number of clonotypes in each sample (X axis) from the validation batch. All numbers are given in terms of unique rearrangements; matching to associated clonotypes allows up to one amino acid substitution in CDR3. **C, D**. Distribution of COVID-19-associated TCRα (**C**) and TCRβ (**D**) clonotype fraction across healthy and convalescent samples. **E, F**. CDR3 sequence similarity graph of COVID-19-associated TCRα (**E**) and TCRβ (**F**) clonotypes, where edges (not shown) connect sequences with up to one amino acid substitution. Each connected cluster is highlighted with its own color. Predicted antigen specificity according to VDJdb is shown with arrows and labels. Green labels correspond to SARS-CoV-2 epitopes, red labels to the other viruses. The detailed list of associations is available in **Supplementary Table 1** (TCRα) **and 2** (TCRβ). **G**. Co-occurrence of CDR3 sequences in α and β chain clonotype clusters. Co-occurrence was calculated as the Tanimoto index for the corresponding TCR α and β clusters. The color corresponds to the correlation coefficient between the usage of α and β cluster clonotypes in the specified clusters. Bold squares show α-β pairings that demonstrate significant association with the same antigen according to VDJdb. **H, I**. CDR3 sequence logos of top four largest clusters in **E** and **F** (TCR α and β respectively).

### Assessment of COVID-19-associated TCR biomarkers

We explored our list of COVID-19-associated TCR clonotypes to see if there are some characteristic motifs that may be linked to antigen specificity. Firstly, we calculated the CDR3 amino acid sequence similarity graph with a threshold Hamming distance of 1 for edges (**Figure 3E, F**). The presence of prominent hubs in the graph suggests that the clustering is non-random, and that there may be large groups among the identified biomarkers that share the same CDR3 motifs and have similar antigen specificity. After computing connected components, we defined 115 and 58 TCR α and β clonotype clusters, or metaclonotypes—an order of magnitude reduction in the number of biomarkers compared to the original clonotype list. We repeated the analysis with the validation batch dropped which also resulted in a sufficient clustering (**Supplementary Figure 6E, F**).

We then performed a search against the VDJdb database^32^ to query for potential antigen specificities of metaclonotypes, allowing for a single amino acid substitution in the CDR3 region. In order to find reliable specificity predictions, we tested if the fraction of clonotypes associated with a given antigen in our cluster is higher than the fraction observed for the entire clonotype list. In cases when multiple antigens are associated with the same cluster, we selected the most enriched one. After setting a P < 0.05 threshold for the Fisher exact test and choosing the epitopes with maximum enrichment score, we found that 23 and five TCR α and β clusters, respectively, were strongly associated with SARS-CoV-2 epitopes (**Supplementary Table 1** and **2**). Three matches to epitopes other than SARS-CoV-2 were found for β clusters, with 27 matches to distinct epitopes for α clusters. The latter might be either false-positive hits, arising as a consequence of us considering only one chain whereas specificity is defined by both. Alternately, they may represent bystander clonal expansions in COVID-19 due to structural relatedness between the original epitope and SARS-CoV-2-derived antigens—for example, the association of α cluster#3 with an epitope from HCoV-HKU1.

In order to overcome the limitation of the absence of TCR α and β pairing information, we performed an *in silico* chain pairing procedure by studying the correlation of α and β clonotypes across samples. The detailed description of the approach applied can be found in **Methods** section. This approach has limitations, as it will not be able to detect pairing for extremely rare or extremely frequent metaclonotypes. The resulting heatmap of α and β pairing for our clusters is shown in **Figure 3G**. We next performed antigen specificity prediction for pairs of clusters, using only those VDJdb records that contain both TCR chains. Interestingly, only strongly-correlated TCRα and β clusters matched to the same epitope according to our annotation, and many of these epitopes were associated with SARS-CoV-2 (**Supplementary Table 3**), although there were also some matches to Epstein-Barr virus epitopes and even neoantigens that may be attributable to cross-reactivity. Characteristic CDR3 amino acid motifs for the four largest α and β clusters are shown in **Figure 3H** and **I**. The full database of biomarkers is available online and can be accessed via https://covidbiomarkers.cdr3.net/.

### Building a COVID-19 status classifier using TCRα and β biomarkers

We subsequently used these TCRα and β biomarkers to build an accurate COVID-19 status classifier. We consider both clustered metaclonotypes and unclustered clonotypes as features, training for either one or both chains. Note that here we considered the biomarker list obtained for all batches in order to provide uniform coverage of repertoires; leaving some batches out decreased classifier recall (data not shown). Among the possible parameters for a classifier is the choice of biomarker frequency encoding and the addition of binary-encoded HLA features. Biomarker frequency can be encoded as Boolean (present or absent), categorical (supported by 0, 1, 2, or 3+ reads) or real (fraction of reads) values. One can also add binary-encoded HLA information, based on the expectation that biomarkers will have different feature importance depending on the HLA context. After training an example support vector machine (SVM) classifier (**Supplementary Figure 7**), we found that real biomarker frequency encoding gives the highest accuracy, with almost no effect from the inclusion of HLA information. The non-informativeness of HLA data can be explained by the fact that these were obtained from the entire dataset, with a diverse mixture of HLAs, and thus had to be robust to donor HLA haplotype. This is further supported by our observations regarding classifier training on allele-restricted set of samples, described later in the text.

As some batches contained exclusively healthy or COVID-19-convalescent individuals, we combined them to balance the number of cases as described in **Methods**. As before, we withheld batch #6, which is relatively balanced, to monitor for batch effects in classifier training. We considered six commonly-used classifier models and measured the F1 score, which is robust to case imbalance, to select the optimal classifier based on training with various sets of features (*e*.*g*., using one or both chains, considering metaclonotype biomarkers). SVM showed optimal performance, with multi-linear perceptron (MLP) having a slightly lower median score (**Figure 4A**), and the former was chosen for in-depth evaluation of classifier performance. As seen in **Figure 4A**, using both α and β chains simultaneously results in a highly accurate classifier, suggesting that biomarkers from each chain convey additional information that benefits overall classification. This is supported by the fact that classification results obtained using α and β chain biomarkers are independent (**Figure 4B**, Pearson R = 0.44). Interestingly, using metaclonotypes in the αβ classifier provides far better separation between COVID-19-convalescent and healthy cases (**Figure 4C**), perhaps owing to less sparsity in feature values. All these factors result in optimal classifier metrics (**Figure 4D, E, F**), leading to excellent sample classification (**Figure 4G**).

**Figure 4.**
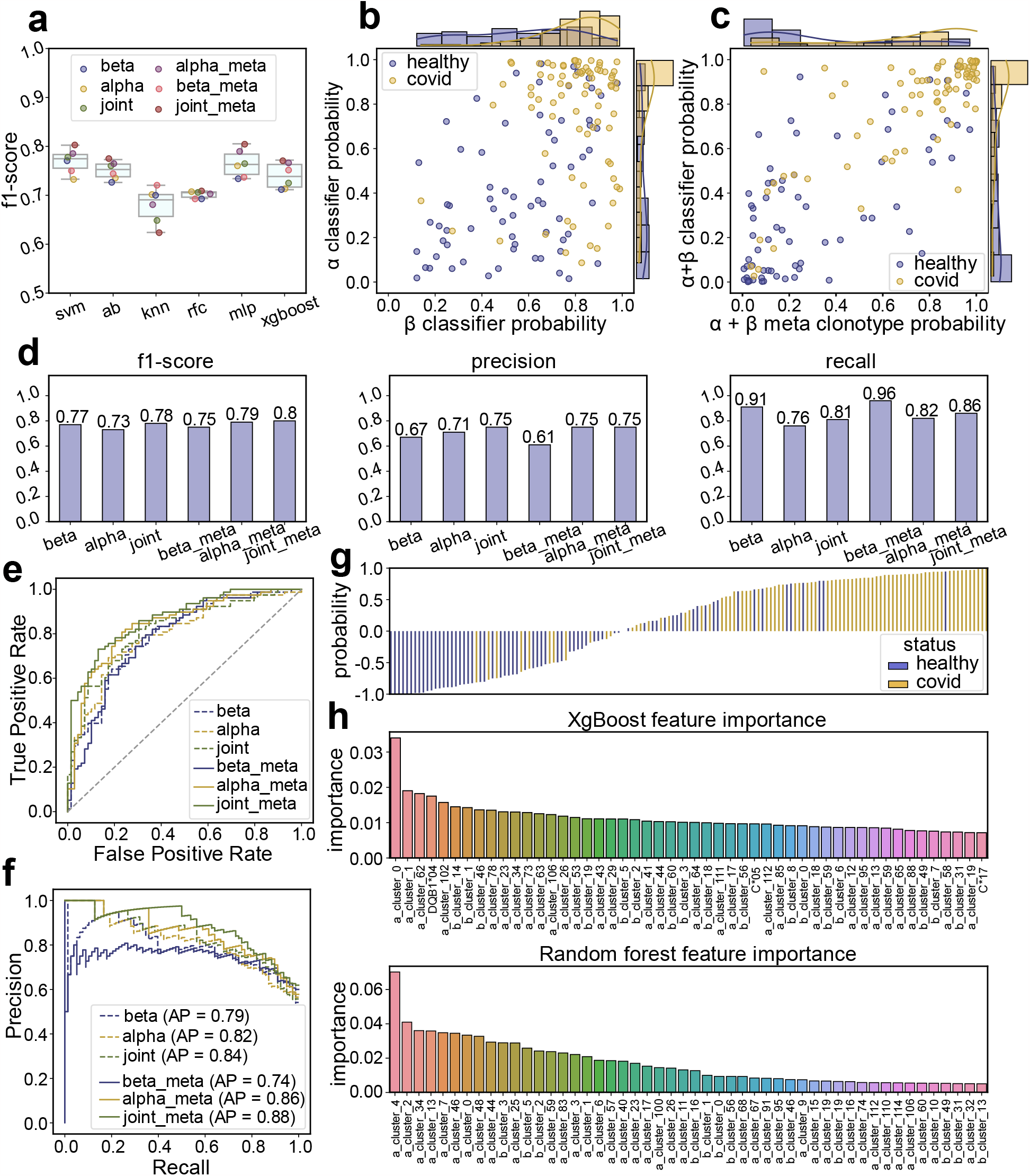
Building a classifier from TCRβ and α biomarkers. Classifiers were trained on all batches except #6, which was left out for validation purposes. **A**. Comparison of ML model F1 scores produced for each feature set. **B**. Scatter plot of the probabilities of labeling samples as COVID-19-positive for TCRα- and β-based classifiers. Histograms on the periphery show the distribution of probabilities for each variable. **C**. Scatter plot of the probabilities of labeling samples COVID-19-positive for classifiers based on both TCRα+β biomarkers and TCRα+β metaclonotype cluster features. The periphery plots are similar to **B. D**. Target metrics (F1-score, precision, recall) for all evaluated models. **E**. Receiver operating curve (ROC) for SVM-based classifiers for all sets of features. **F**. Waterfall plot of the probability of each sample being labeled as COVID-19 positive (> 0) or healthy (< 0). Samples from healthy donors are blue, COVID-19 samples are orange. **G**. Precision-recall curve for SVM-based classifiers using different biomarker sets. **H**. Feature importance plot for XGBoost or RandomForest classifier models based on TCRα- and β-based meta-biomarkers and HLA features.

We additionally assessed the impact of batch effects on classifier performance. We performed a leave-one-batch-out cross-validation procedure to compute classifier accuracy, and focused on the difference between combined batches (batches 4+1, batches 4+2, batches 4+3, batches 4+9), batches 5, 7 and 8, and held-out batch 6 (**Figure S8**). As expected, the combined batches demonstrated a higher score due to batch effects; however, batch #6, which was not used at the biomarker selection step, was classified with accuracy similar to batches 5, 7 and 8, suggesting that our set of biomarkers is robust. We analyzed the feature importance of the TCRα+β metaclonotype-based classification by training XgBoost and RandomForest classifiers (**Figure 4H**). RandomForest almost exclusively ranked α chain features as being most important, with a couple of β chain features showing medium to low importance. XgBoost model, on the other hand, ranked many β chain features as being important (17 out of 58 β metaclonotypes among the top-50 most important features), and additionally included 3 HLA-related features. It may be that XgBoost’s ranking of β and HLA features is due to its ability as a boosting model to highlight features that are important for small subsets of data.

### Applying the COVID-19 status classifier to an independent dataset

We use a previously published data compendium from Adaptive Biotechnologies^8^ (Cohort II) to validate our classifier, and also to see whether we can train an efficient classifier on this cohort and apply it to Cohort I. Cohort II comprises 777 healthy and 1,214 COVID-19 samples prior after coverage filtering **(Supplementary Figure 1C, D)**, and is limited to TCRβ chain data. The healthy donors were taken from the HIP and KECK cohorts, which were initially used for CMV infection status research^25^. The main problem with this dataset is that the healthy and COVID-19 convalescent samples come from different batches sequenced years apart, which can result in strong batch effects. Moreover, this dataset was generated using a proprietary protocol and primers that are different from the ones used here, such that a strong batch effect is also expected when comparing Cohorts I and II. We applied the batch effect correction technique discussed above, and resampled the clonotypes afterwards (**Figure 5A, B**). As the average number of reads (sequencing depth) in Cohort II is far larger than in Cohort I, each sample in this cohort was down-sampled to 50,000 reads. We repeated the feature selection pipeline for the resampled Cohort II data, and found 17,185 TCRβ biomarkers significantly associated with COVID-19 status (compared to ∼500 in Cohort I). However, due to our inability to decouple COVID-19 status and batch, we suspect the majority of these features reflect batch effects. To narrow down the biomarker list, we used a log enrichment (observed:expected incidence ratio in COVID-19 samples) threshold of 2.5, yielding 2,066 biomarkers (**Figure 5C**). Interestingly, there were no spurious clonotypes, and clonotype incidence rates were in good agreement with those predicted by a theoretical model of V(D)J rearrangement.

**Figure 5.**
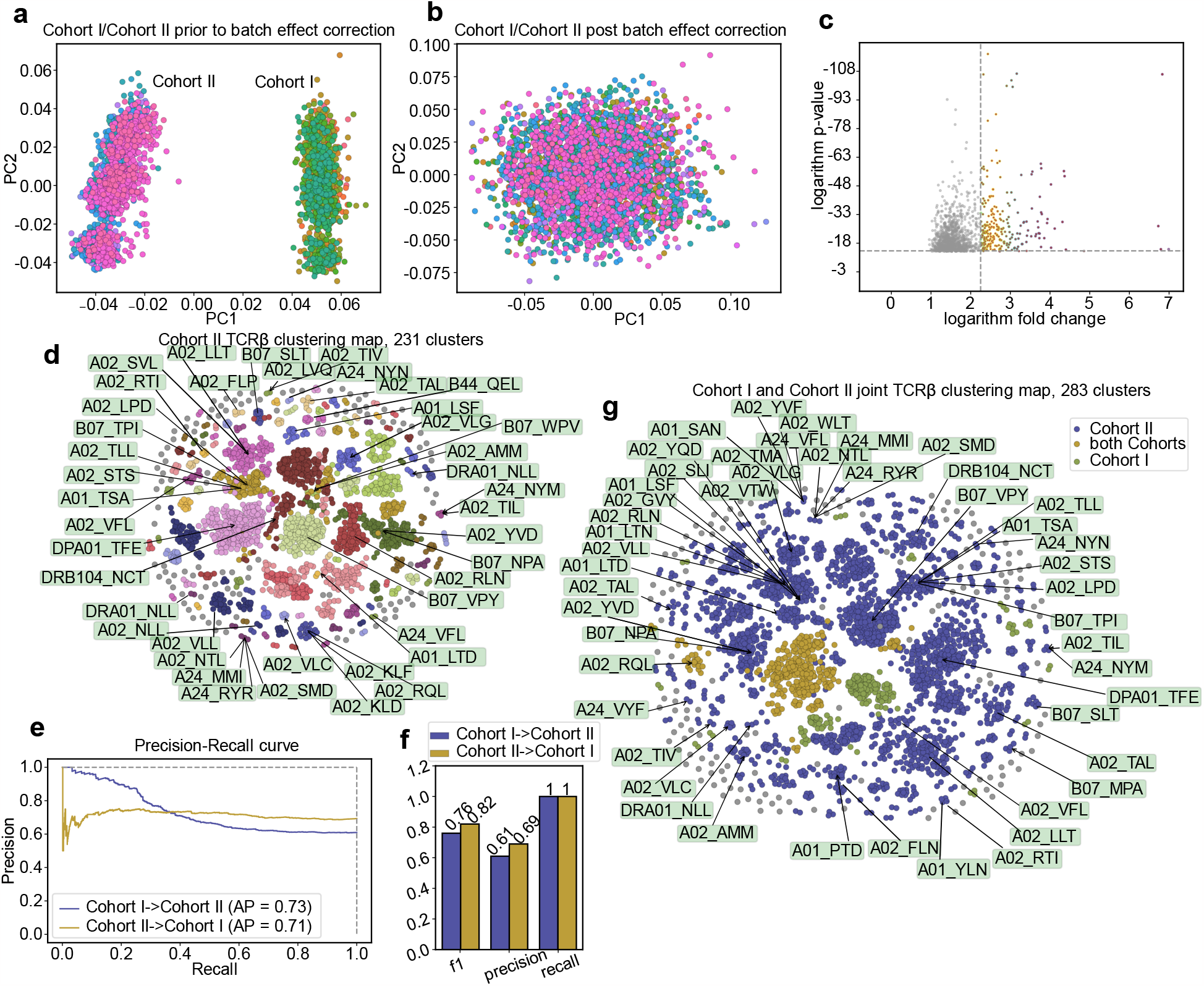
Comparative analysis of classifiers between cohorts. **A, B**. Visualization of batch effects pre-(**A**) and post- (**B**) correction procedure for TRBV genes. Colors show sample batch. **C**. A Fisher test-based approach revealed >20,000 statistically significant clonotypes. The volcano plot was used to select the clonotypes with the highest fold change. **D**. Homology graph of CDR3 sequences of COVID-19-associated TCRβ clonotypes for Cohort II. The most probable epitope is marked for each cluster associated with SARS-CoV-2. **E**. Precision-recall curves for Cohort I (blue)- or Cohort II (yellow)-based classifiers. AP = average precision. **F**. Comparison of metrics for the models described in **E. G**. Homology graph of CDR3 sequences of COVID-19-associated clonotypes for both Cohort I and Cohort II TCRβ biomarkers. Clusters containing clonotypes from both classifiers are colored orange.

We proceeded with similarity clustering of TCRβ clonotypes and arrived at 231 metaclonotypes, 33 of which were specific to SARS-CoV-2 epitopes according to VDJdb (**Figure 5D**). The fraction of SARS-CoV-2 epitope-specific metaclonotypes in Cohort II is 14% (33 out of 231), which is higher than the 8.6% (5 out of 58) observed for TCRβ in Cohort I. We next tested the performance of a classifier trained for Cohort I on Cohort II data and vice versa. First, we took an existing TCRβ classifier trained for 5,670 biomarkers from the Cohort I data and applied it to Cohort II, with prior batch-effect correction between cohorts. In this case, we did not downsample the Cohort II data, as we expected that the Cohort I biomarkers would be quite rare and require additional depth to be detected in Cohort II. Unfortunately, 15 of the described biomarkers were still absent in all samples from Cohort II. This approach resulted in a validation F1-score of 0.76, precision score of 0.61, and recall of 1 for the preselected model, and an optimal precision-recall (PR) curve (**Figure 5E, F**). We also trained an SVM classifier on Cohort II biomarkers, downsampled the data, and applied it to Cohort I. In this case, we got roughly similar performance results, but with suboptimal average precision (**Figure 5E, F**). It may be that the design of Cohort II, where COVID-19 status cannot be decoupled from batch effects, does not allow training an optimal classifier. Nevertheless, when we checked whether TCRβ clonotype biomarkers identified separately for Cohort I and II have similar sequences, we observed five overlapping clusters of clonotypes—strong evidence that these two classifiers homed in on similar clonotypes (**Figure 5G**).

### HLA association analysis reveals a unique COVID-19 biomarker motif

Imprints of past and present infections on TCR repertoires are associated with individual HLA haplotypes, as the set of antigens available to T cells is shaped by HLA restriction^33^. The same should be the case for COVID-19, and so we searched for TCRs associated with both donor status and HLA haplotype, employing a strategy proposed previously^28^. HLA-associated clonotypes were found by using the Fisher exact test to compare incidence in individuals with a specific HLA allele versus those without it. We then calculated COVID-19 association for the subset of samples having an HLA of interest. After obtaining lists of HLA-associated clonotypes and COVID-19-associated clonotypes for every HLA, we identified their intersection in order to generate a list of HLA-restricted biomarkers. This allowed us to explore rare clonotypes present in COVID-19 patients depending on HLA context. This approach led to the identification of 13 TCRβ biomarkers that were exclusive to COVID-19 donors with HLA-DQB1*05 or HLA-DRB1*16 alleles which are present in COVID-19 positive samples from all the batches (**Supplementary Figure 9A**). Interestingly, while the search was performed using a CDR3 region that contains only a few amino acids (*i*.*e*. a CASS subsequence that is common across many TRBV alleles), all of these clonotypes were recombinants that incorporated TRBV12-3. The amino acid motif corresponding to the 13 clonotypes generally featured only one variable position (position 8), whereas all other positions remained largely fixed (**Figure 6A**). It is also interesting to note that the presence of both HLA-DQB1*05 and HLA-DRB1*16 (which have high linkage disequilibrium) is associated with various autoimmune diseases^34,35^. This pattern of clonotypes was found in 75 donors and was almost always associated with positive COVID-19 status (63 out of 75 donors; **Figure 6B**), allowing straightforward classification of a fraction of convalescent donors with a common HLA allele. Moreover, 11 of the 12 healthy donors with clonotypes matching the found motif had only one read representing the clonotypes of interest; as such, when we dropped all samples having just one read with the corresponding pattern, the HLA linkage became even more evident (**Figure 6C, Supplementary Figure 9B**).

**Figure 6.**
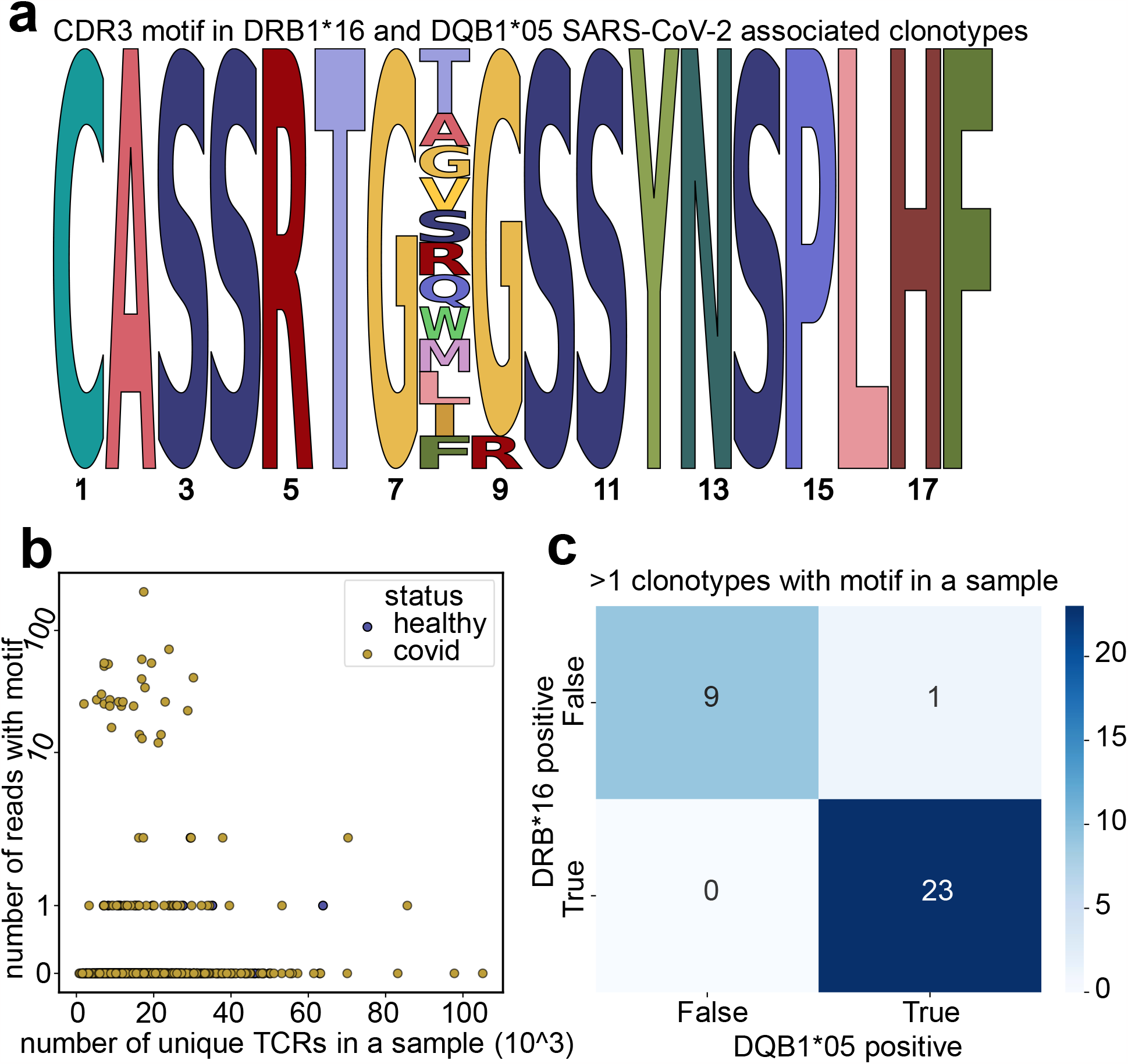
Analysis of DRB1*16, DQB1*05 and COVID-19-associated TCRβ biomarkers. **A**. Consensus motif for the 13 TCRβ biomarker associated with the DRB1\**16, DQB1\**05 HLA alleles and COVID-19 history. **B**. The number of TCRβ motif biomarker reads versus number of unique TCR sequences in healthy and COVID-19 cohorts. **C**. Heatmap representing the linkage of HLAs of interest for all the COVID-19 patients with at least two motif reads present in a sample.

### Separate HLA classifiers provide better classification quality

Although our previous attempts to incorporate HLA information into the COVID-19 classifier didn’t yield any gains in accuracy, we hypothesized that this might be due to limited cohort size and the presence of a large number of rare HLA alleles. Thus, we decided to directly check whether HLA information can improve classifier performance by selecting samples carrying a certain HLA allele for our pipeline. We chose HLA-A*02^+^, as this is among the most common HLA alleles—in Cohort I, there were 545 positive donors for TCRα and 521 positive donors for TCRβ. We then repeated our COVID-19 association analysis, identifying 3,277 and 358 TCRα and β clonotype biomarkers that form large hubs in a TCR similarity graph, and which can respectively be clustered into 174 and 197 TCRα and β metaclonotypes (**Figure 7A, B)**. The majority of these were specific to HLA-A*02-restricted epitopes, according to VDJdb annotation. Next, we used this HLA-A*02-restricted biomarker set to train classifiers for the HLA-A*02^+^ subset of the cohort, resulting in highly accurate COVID-19 status predictions for SVM classifier (**Figure 7C, D**). To perform a proper comparison with the HLA-unrestricted classifier, we repeated the whole procedure described in this section for 545 and 521 Cohort I donors for TCRα and β chains, respectively, who were randomly selected regardless of their HLA. The resulting classifier performed worse than the classifiers for our entire cohort, as we had decreased the dataset size by nearly two-fold. These results demonstrate that HLA information can greatly boost classifier performance when one is provided with a sufficient number of samples with a given HLA and is handling them correctly. Moreover, our approach can be utilized for accurate HLA allele prediction for frequent alleles (**Supplementary Figure 10**).

**Figure 7.**
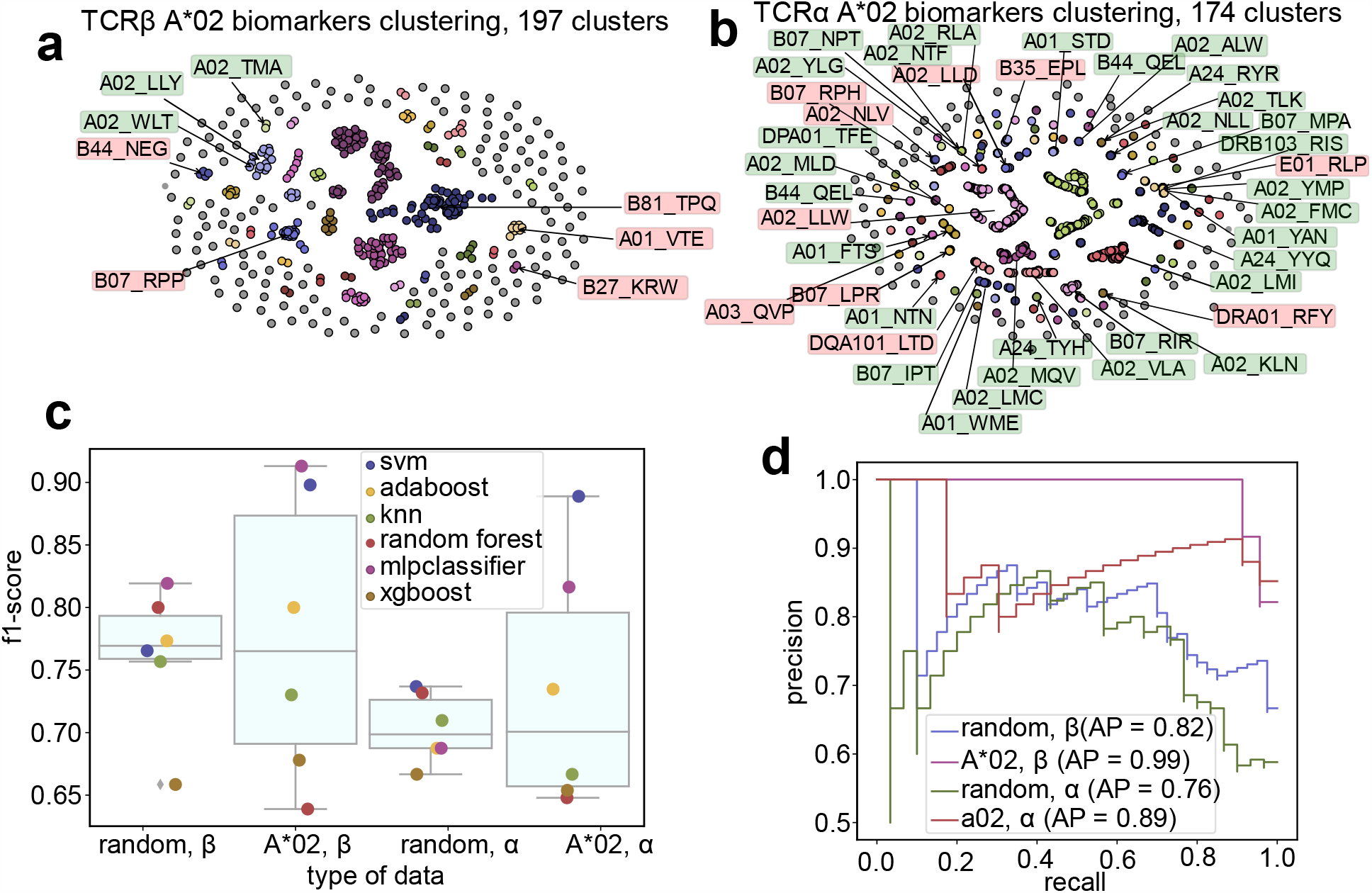
Comparison of classifiers built using biomarkers derived from random samples and A*02^+^ samples. **A**. Homology graph of CDR3 sequences of COVID-19-associated TCRβ clonotypes derived using only A*02^**+**^ samples. For each cluster associated with SARS-CoV-2, the most probable epitope is marked. Green labels show SARS-CoV-2 epitopes, red labels belong to other viruses. Detailed information on cluster associations is available in **Supplementary Data 5. B**. Same as **A**, but for TCRα biomarkers. Only matches for clusters of size >9 are shown. For more information see **Supplementary Data 5. C**. Comparison of model performance for different datasets. Random alpha and beta datasets correspond to classifiers built using biomarkers derived from 545 and 521 random TCRα and β samples, respectively, in Cohort I. A02 datasets correspond to classifiers built for A*02^+^ biomarkers and samples. **D**. Precision-recall curve for SVM classifiers built for the datasets in **C**. Classifiers using single HLA allele samples and single HLA allele SARS-CoV-2-associated clonotypes provided better classification.

### Classification based on published SARS-CoV-2 epitope-specific TCR sequences

Our previous results indicate that many inferred biomarkers are similar to TCR sequence variants that recognize SARS-CoV-2 epitopes according to previously published studies. Thus, we decided to test whether we can create an accurate COVID-19 status classifier by solely relying on publicly available TCR sequences as biomarkers. We started with 2,909 and 3,816 SARS-CoV-2 epitope-specific TCRα and β sequences, respectively, from the VDJdb database^32^ as a biomarker set. Overall, there was no significant difference in the incidence of clonotypes matching VDJdb biomarkers (with up to one amino acid substitution) in convalescent donors compared to healthy ones. We decided to narrow the set and proceed with Fisher test analysis for COVID-19 status association. We observed no significant difference for the majority of clonotypes, as most of them are found in only a handful of samples in the dataset. Nevertheless, we were able to select 338 TCRβ (**Supplementary Figure 11A, B, C**) and 265 TCRα (**Supplementary Figure 11D, E, F**) biomarkers for further analysis. We evaluated classifiers based on this set of biomarkers and found that it yields an F1 of 0.6–0.7, which is substantially lower than the classifier built using Cohort I data (F1 = 0.7–0.9; **Supplementary Figure 11G–J**). However, one can still see the improvement of classification metrics in the paired-chain classifier model compared to the single-chain one. These results suggest that the number of publicly-available TCR specificity records is still too small to enable accurate COVID-19 status prediction in a large and diverse population of donors.

## Discussion

Since the advent of AIRR-seq technology, there has always been an expectation that it will yield a wealth of novel diagnostic biomarkers and complement commonly-used immunological assays in infectious disease and vaccine studies. However, the complexity of the AIRR-seq protocol—combined with the extreme diversity of antigen receptors, variability in the strength of immune response in different donors, and the phenomenon of public (shared across multiple donors) and private immune responses—make it extremely hard to employ this technique in a clinical setting. In the present study, we used a large cohort of COVID-19-convalescent and healthy donors to design a proof-of-concept wet lab and bioinformatic pipeline based on AIRR-seq data that can be employed to accurately stratify patients based on the history of infection.

The main obstacle in the way of designing a universal pipeline based on a robust set of COVID-19 TCR biomarkers is the presence of strong batch effects in the AIRR-seq data that can lead to data overfitting and misclassification of donor samples^23^. As observed previously, most batch effects manifest in the form of skewed V and J gene usage profiles and the presence of ultra-public TCR variants due to cross-sample contamination. Here, we designed and applied a gene-usage normalization strategy that was able to remove biases between batches and technologies, making our data consistent and comparable with previously published datasets. This approach is useful in cases where each donor group is sequenced in a separate batch^8^, and can more broadly benefit research comparing AIRR-seq data from different groups of donors or entailing meta-analysis of different AIRR-seq studies^41^. We also propose the use if V(D)J rearrangement model software such as OLGA^31^ to control for potentially spurious clonotypes arising from artifacts or contaminants. Importantly, we explicitly show that our approach can derive a set of biomarkers that can be transferred and applied to an independent COVID-19 dataset obtained from a population with distinct demographics using a different AIRR-seq protocol.

The ability to sequence only one TCR chain is one of the major limitations of bulk AIRR-seq methods. Most studies limit their analysis to TCRβ sequencing data^42^, even though interaction with both chains is required for proper antigen binding^43^ and there are multiple cases of prominent TCRα motifs recognizing immunogenic epitopes^41,44^. In the present study, we clearly demonstrate that TCRα and β chain biomarkers contain complementary information and that joint analysis of both chains can greatly boost COVID-19 status classifier performance even in the absence of single-cell-resolution pairing data^40^. The latter can be partially recovered by performing *in silico* pairing based on biomarker co-occurrence.

Given that T-cells observe pathogens through the prism of HLA alleles presenting distinct peptides, we combined the AIRR-seq data reported in this paper with donor HLA typing. Our results show that in general, unsupervised feature selection leads to an HLA-agnostic set of biomarkers, and that classifiers built using these cannot be improved by including donor HLA information. On the other hand, dedicated classifiers trained on donors with shared HLA alleles (*e*.*g*., the HLA-A*02 allele, which is relatively common in the population) showed higher accuracy than those trained for sets of donors of similar size with mixed HLA alleles or for the whole dataset. We were also able to find a relatively rare (in terms of incidence in donors) subset of biomarkers that is nevertheless tightly linked to donor HLA class II haplotype DRB1*16/DQB1*05 and COVID-19 positive status. Overall, our findings suggest that HLA information can greatly benefit AIRR-based donor classifiers if handled properly^45,46^.

The pipeline described in the present study can be easily adapted to AIRR-seq studies of various infectious diseases and pathologies. Furthermore, the set of biomarkers produced in this work can be applied to monitor T cell memory for COVID-19 in donor samples and test if similar TCR sequences are induced by vaccination with SARS-CoV-2 antigens. We hope that further AIRR-seq studies containing large and diverse cohorts of donors will extend the TCR biomarker knowledgebase to other conditions, and make AIRR-based diagnostics and monitoring more common in clinical settings in the near future.

## Acknowledgements

This work was supported by a grant from the Ministry of Science and Higher Education of the Russian Federation (075-15-2019-1789). Authors are grateful to Michael Eisenstein for the helpful edits.

## Materials and Methods

### Cohort collection

This study is a multicenter cross-sectional stratified epidemiological research. The study was conducted in Federal Medical-Biological Agency (FMBA) medical organizations in 54 regions of the Russian Federation. The design of the study included volunteers from two cohorts. The first, ‘control’ cohort included conditionally healthy people without signs of respiratory diseases at the time of inclusion in the study and a SARS-CoV-2 PCR negative test, as well as samples of healthy individuals collected before the COVID-19 pandemic. The second ‘core’ cohort included people with COVID-19, with a positive test for SARS-CoV-2, lung CT findings, and a history of diagnosis. This core group included participants with different severities of COVID-19—asymptomatic, mild, moderate, severe, or extremely severe—as defined in the recommendations of the Ministry of Health of the Russian Federation. The core group included patients who had gone no more than 28 days since the diagnosis of COVID-19. A total of 1,253 participants were included in this study (392 in the control group and 861 in the core group) in the period from June to October 2020. All participants underwent a medical inspection, physical examination, collection of blood samples, and oropharyngeal swab. Samples were stored at 4–6 °C for no more than two hours after sampling and transfer to the laboratory, and were consistently maintained at -80 °C until time of analysis.

### Ethical considerations

The study was conducted according to the guidelines of the Declaration of Helsinki, and approved by the Local Ethics Committee of the Federal State Budgetary Institution “Centre for Strategic Planning and Management of Biomedical Health Risks” of the FMBA (Protocol No. 2 of May 28, 2020). All participants provided informed consent.

### DNA isolation

For HLA typing, genomic DNA (gDNA) from whole blood samples was isolated manually using the DNA Blood Mini Kit (Qiagen, Germany) in accordance to the manufacturer’s protocol. The yield and purity of the isolated gDNA were manually determined using a Qubit 4.0 fluorimeter (Thermo Fisher Scientific, USA) and NanoDrop One C Microvolume UV-Vis (Thermo Fisher Scientific, USA), respectively. Only gDNA samples with absorbance ratios of 1.7–1.9 A_260/280_ and 1.8–2.2 A_230/260_ were selected for further analysis. For TCR library preparation, 200 μl of frozen whole blood was used for gDNA isolation using the Amplitech E1 automatic sample preparation station (LLC “Amplitech”, Russia). The first step was lysis of blood samples, followed by washing. The gDNA was eluted with 50 μl of TE buffer, and the gDNA concentration was manually determined using a Qubit 2.0 fluorimeter (Thermo Fisher Scientific, USA).

### HLA typing

A total of 150-500 ng of gDNA was used to prepare high-throughput sequencing (HTS) libraries. Libraries were prepared using the Illumina DNA Prep kit (Illumina, Inc. USA) according to the manufacturer’s recommendations with a Tecan Freedom EVO robotic station (Tecan, Switzerland). The size of the resulting libraries was determined with the Agilent D1000 reagent kit and Agilent 4200 TapeStation (Agilent Technologies, USA). Pooling was performed automatically using a Tecan Freedom EVO robotic station (Tecan, Switzerland). Pool quality control was performed with an Agilent High Sensitivity D1000 Screen Tape reagent kit using the Agilent 4200 TapeStation (Agilent Technologies, USA). Whole-genome sequencing (WGS) was performed using the Illumina NovaSeq 6000 with the S4 reagent kit (300 cycles) (Illumina, USA) with 2 × 150 bp paired-end reads. Samples were typed at six main loci of HLA classes I and II (HLA-A, -B, -C, -DRB1, -DPB1, -DQB1) with a resolution of two fields with the xHLA bioinformatic software^36^.

### TCR library preparation

The sequencing libraries for TCR α and β chains profiling were prepared using Human TCR DNA Multiplex kit (MiLaboratories, USA) according to the manufacturer’s protocol. Briefly, 600 ng DNA (equivalent to 92,000 genome copies with expected ∼50% related to T cells) extracted from whole blood was used as input in two separate 50 μl multiplex PCR replicates containing 1X Turbo buffer, 10 U HS Taq polymerase, 200 μM of each dNTP (all Evrogen, Russia) and 1X TR hum DNA PCR1 primer mix from the Human TCR DNA Multiplex kit. The amplification profile was 94 °C for 3 min, followed by 8 cycles of 94 °C for 20 s, 57 °C for 90 s, and 72 °C for 40 s, followed by an additional 17 cycles of 94 °C for 20 s and 72 °C for 80 s. Denaturation steps (94 °C) were performed with a ramp of 4 °C/s, and other steps were performed with a ramp of 0.5 °C/s. 10 μl of each replicate amplicon were pooled and purified using 1V AmPure XP beads (Beckman Coulter) and used as a template for the second PCR. The second (indexing) 25 μl PCR contained 1X Turbo buffer, 2.5 units of HS Taq polymerase, 200 μM of each dNTP, and 1 μl of unique dual-index primers (Illumina, USA). The amplification profile was as follows: 94 °C for 3 min, followed by 15 cycles of 94 °C for 20 s, 55 °C for 20 s, and 72 °C for 40 s. Obtained amplicons were purified using 0.8V AmPure XP beads (Beckman Coulter, USA), pooled, and sequenced on the Illumina NovaSeq 6000 system in 150PE mode.

### External datasets

Cohort II data from Adaptive Biotechnologies was obtained from Emerson *et al*.^25^ (HIP and KECK batches, pre-COVID-19 control) and Nolan *et al*.^8^ (convalescent donor data).

Data processing and analysis

All datasets were processed with MIGEC and MiXCR software as reported previously^27,37^. Clonotype sequences for Cohort II datasets were re-mapped using MiXCR to keep consistent V(D)J naming across datasets. Clonotype tables were formatted and standardized using VDJtools software ^38^. Formatted datasets can be downloaded from Zenodo^39^, and the list of TCR biomarkers can be downloaded at https://covidbiomarkers.cdr3.net/. All data analysis was performed using in-house R and python scripts available at GitHub (https://github.com/antigenomics/tcr-covid-classifier), and the analysis can be reproduced by running a Snakemake workflow stored in the repository.

### Batch-effect correction and data normalization

We focused on sample preparation and sequencing biases manifested in the difference in frequency of V and J gene rearrangements across batches and normalized samples as follows. We assumed that gene usage (*P*, fraction of reads mapping to a given gene in a given sample) is log-normally distributed for each gene across samples in a given batch, with a specific mean and standard deviation *P(gene, sample) ∼ LogNormal(μ, σ* | *batch)*. We therefore log-transformed gene usage and computed *Z*-scores, where *Z =* (*log P - μ*) / *σ* for each gene, so that the distribution of gene usage becomes a standard normal distribution *N(0, 1)* across all samples. As we performed the analyses at the clonotype level, we needed the means to re-define and correct raw clonotype frequencies (*f*_*i*_), and we therefore put the *Z*-scores back into the *[0, 1]* interval so that we can use them to scale clonotype frequencies. We used sigmoid transform for this purpose, adjusted to preserve the original average gene frequency *P*_*avg*_*(gene)* across all batches. We then computed target frequencies as *P*_*final*_*(gene, sample) = 2* × *P*_*avg*_*(gene)* / *(1 + exp[-Z(gene, sample)])*. We can then correct the frequencies of clonotype *f*_*i*_ from sample *S* rearranged from gene *G* as *f*_*i*_*’ = f*_*i*_ *× P*_*final*_*(G, S) / P(G, S)*. We next adjusted the sample clonotype composition according to the new frequencies. As our frequency adjustment can lead to non-integer clonotype counts, straightforward sampling from a multinomial distribution is not feasible. We therefore randomly sampled numbers from a uniform distribution *U[0, 1]* equal to the read count in the sample and performed a roulette-wheel selection for clonotypes according to their adjusted frequencies *f*_*i*_*’’* normalized to a sum of *1*.

### *In-silico* α-β pairing of clonotype groups

This approach is inspired by the work of Howie *et al*.^40^ and extended by the addition of fuzzy CDR3 matching and working with clonotype groups instead of clonotypes for robustness. We estimate the pairing between selected TCRα and β chain clonotype groups as follows. For a given group of clonotypes *g*, we estimated the number of matching clonotypes in a given sample *s* as *X*_*gs*_, which is equal to the number of CDR3 amino acid sequences in the sample that match at least one sequence in *g* with up to one amino acid mismatch allowed (*i*.*e*. Hamming distance ≤ 1). As an example, consider the i^th^ group of α chain clonotypes and j^th^ group of β chain clonotypes. We can compute the Pearson correlation, ρ_*ij*_ , of X_is_ and X_js_ across all samples and use it as a score for pairing between these two groups. Alternatively, one can compute *n*_*i*_ = ∑_*s*_ *X*_*is*_ > *0* and *n*_*ij*_ = ∑_*s*_ *X*_*is*_ > *0* ∧ *X*_*js*_ > *0* and define the odds ratio of detecting a given pair as θ_*ij*_ = (*n*_*ij*_ *× n*) */* (*n*_*i*_ *× n*_*j*_), where n is the total number of samples. Note that the potential of *in silico* pairing is limited. A too-frequent group with high incidence across samples will be paired with almost all groups of another chain, whereas a rare group may have insufficient statistical power to detect a matching group.

### Clonotype group antigen specificity annotation

We annotated clonotype groups with the putative antigen they recognize by matching TCRα and β sequences to antigen specificity records from the VDJdb database^32^. In general, we considered a match between CDR3 sequences with no more than a single amino acid substitution relative to a database record to indicate that a given TCR sequence recognizes a corresponding antigen. In this way, we arrived at the putative set of antigens *{A}*_*G*_ for a given clonotype group *G*. The overall matching between a given clonotype group and antigen *A* can be assessed as follows: let *n*_*A*_ be the number of matching clonotypes in a group of interest, *m* be the size of group of interest, *N*_*A*_ be the number of matches in all groups and *M* be the total number of clonotypes in groups. We can then use binomial distribution with parameters *n*_*A*_, *m* and *p* = *N*_*A*_ */ M* to test the significance of overlap *n*_*A*_ */ m* as described in Pogorelyy *et al*.^28^.

For a given α and β chain clonotype group pair *(Gα, Gβ)*, we can narrow down the search to the intersection of their putative antigen sets. We can exploit the true α and β pairing information present in VDJdb to find high-confidence antigen matches for pairs of clonotype groups as follows. Let us consider only paired records in VDJdb for an antigen *A*; we next iterate over all pairs of TCRα and β sequences in our clonotype group and see if they match any paired database records with a single amino acid substitution in both CDR3 chains allowed. Counting the extent of pairing *x*_*A*_ (*i*.*e*., the total number of TCRα and β sequences matched to *A*), we can compare it to the expected value by using the hypergeometric test with the maximum potential number of matches *X*_*A*_ = *min(*|*Gα*|, |*Gβ*|), the number of paired records belonging to antigen *A* in VDJdb, and the total number of paired records in VDJdb.

### Selecting a COVID-19 classifier model

We compared a set of commonly used machine learning approaches in terms of their ability to classify COVID-19 status using TCR repertoire data, a set of pre-selected TCR features, and HLA haplotype. These included SVM, AdaBoost (*ab*), KNN, Random Forest Classifier (*rfc*), Multi Linear Perceptron Classifier (*mlpclassifier*), and XgBoost. These models were compared according to their performance, and *ab* and XgBoost were additionally used to analyze feature importance. Grid search based on five-fold cross-validation was used to optimize the parameters of each classifier model for training data that includes all batches except #6, which was withheld from classifier model selection and training, and used for validation.

As the data was still imbalanced, we focused on the F1-score (harmonic mean of the precision and recall metrics) and precision-recall (PR)-curves to assess classifier performance. The best model was chosen by the maximum F1-score on the test sets. We obtained final metrics for best models using leave-one-batch-out cross-validation. Since batches 1–3 contained no negative cases and almost all donors in batch 4 were positive, we constructed merged batches batch14, batch24 and batch34 by evenly distributing batch 4 to the first three batches, and used these instead of original batches 1-4.

