## Supplementary Materials for "Robust detection of SARS-CoV-2 exposure in the population using T-cell repertoire profiling"

#### Supplementary table 1. Epitopes associated with TCRα clusters for Cohort I.

| antigen.epitope | antigen.species | count | pval | cluster | all_clust_count | enrichment_score |
|---|---|---|---|---|---|---|
| PQPELPYPQPE | Wheat | 13 | 0.001307 | 0 | 13 | 1 |
| LMIERFVSL | SARS-CoV-2 | 20 | 5.01E-21 | 0 | 20 | 1 |
| LTDEMIAQY | SARS-CoV-2 | 22 | 8.44E-08 | 0 | 66 | 0.333333333 |
| AYAQKIFKI | CMV | 31 | 1.23E-07 | 0 | 32 | 0.96875 |
| MEVTPSGTWL | SARS-CoV-2 | 33 | 0.000115 | 0 | 34 | 0.970588235 |
| IVTDFSVIK | EBV | 52 | 1.26E-05 | 0 | 211 | 0.246445498 |
| SPRWYFYYL | SARS-CoV-2 | 52 | 3.12E-05 | 0 | 281 | 0.185053381 |
| LLWNGPMAV | YFV | 60 | 2.88E-17 | 0 | 138 | 0.434782609 |
| NLVPMVATV | CMV | 92 | 1.37E-17 | 0 | 642 | 0.143302181 |
| GILGFVFTL | InfluenzaA | 93 | 4.39E-11 | 0 | 504 | 0.18452381 |
| QPRAPIRPI | EBV | 6 | 4.06E-07 | 1 | 6 | 1 |
| MFLARGIVF | SARS-CoV-2 | 29 | 1.61E-07 | 1 | 29 | 1 |
| QYIKWPWYI | SARS-CoV-2 | 35 | 0.001283 | 1 | 68 | 0.514705882 |
| SGPLKAEIAQRLED | InfluenzaA | 36 | 2.03E-08 | 1 | 54 | 0.666666667 |
| KTFPPTEPK | SARS-CoV-2 | 43 | 0.000153 | 1 | 64 | 0.671875 |
| LLWNGPMAV | YFV | 53 | 1.89E-06 | 1 | 138 | 0.384057971 |
| RLRAEAQVK | EBV | 71 | 0.00087 | 1 | 198 | 0.358585859 |
| IVTDFSVIK | EBV | 75 | 7.13E-06 | 1 | 211 | 0.355450237 |
| RAKFKQLL | EBV | 106 | 0.000981 | 1 | 239 | 0.443514644 |
| PKYVKQNTLKLAT | InfluenzaA | 47 | 0.000552 | 2 | 78 | 0.602564103 |
| LPRWYFYYL | HCoV-HKU1 | 32 | 9.06E-38 | 3 | 32 | 1 |
| FLRGRAYGL | EBV | 3 | 0.002733 | 4 | 7 | 0.428571429 |
| QVPLRPMTYK | HIV-1 | 12 | 4.40E-08 | 4 | 15 | 0.8 |
| NEGVKAAW | CMV | 19 | 3.87E-09 | 4 | 51 | 0.37254902 |
| PKYVKQNTLKLAT | InfluenzaA | 19 | 2.12E-09 | 4 | 78 | 0.243589744 |
| TTDPSFLGRY | SARS-CoV-2 | 22 | 1.57E-09 | 4 | 87 | 0.252873563 |
| RTLNAWVKV | HIV-1 | 7 | 6.53E-26 | 5 | 9 | 0.777777778 |
| PKYVKQNTLKLAT | InfluenzaA | 8 | 6.57E-30 | 5 | 78 | 0.102564103 |
| IVTDFSVIK | EBV | 13 | 4.41E-07 | 5 | 211 | 0.061611374 |
| GILGFVFTL | InfluenzaA | 33 | 2.06E-11 | 5 | 504 | 0.06547619 |
| KLGGALQAK | CMV | 36 | 1.75E-08 | 5 | 2190 | 0.016438356 |
| RLITGRLQSL | SARS-CoV-2 | 9 | 6.27E-06 | 6 | 9 | 1 |
| YINVFAFPF | SARS-CoV-2 | 13 | 2.78E-13 | 6 | 13 | 1 |
| QVPLRPMTYK | HIV-1 | 3 | 1.90E-17 | 9 | 15 | 0.2 |
| RLRAEAQVK | EBV | 19 | 3.91E-18 | 9 | 198 | 0.095959596 |
| KLGGALQAK | CMV | 21 | 7.24E-05 | 9 | 2190 | 0.009589041 |
| NLVPMVATV | CMV | 21 | 1.23E-23 | 9 | 642 | 0.03271028 |

| | | | | | | |
|---|---|---|---|---|---|---|
| SPRWYFYYL | SARS-CoV-2 | 21 | 1.78E-25 | 9 | 281 | 0.074733096 |
| DPFRLLQNSQVFS | InfluenzaA | 2 | 1.20E-08 | 10 | 2 | 1 |
| APFSEQEQPVLG | TriticumAestivum | 13 | 0.003376 | 10 | 13 | 1 |
| TTDPSFLGRY | SARS-CoV-2 | 13 | 9.95E-12 | 10 | 87 | 0.149425287 |
| HWFVTQRNFYEPQII | SARS-CoV-2 | 23 | 3.80E-15 | 10 | 23 | 1 |
| RISNCVADYSVLYNS | SARS-CoV-2 | 24 | 1.77E-42 | 10 | 24 | 1 |
| KLGGALQAK | CMV | 89 | 5.85E-07 | 10 | 2190 | 0.040639269 |
| ALNTLVKQL | SARS-CoV-2 | 2 | 6.21E-07 | 13 | 2 | 1 |
| RLRAEAQVK | EBV | 2 | 0.002061 | 13 | 198 | 0.01010101 |
| NTFSSTFNV | SARS-CoV-2 | 5 | 1.84E-14 | 14 | 5 | 1 |
| PFPQPELPY | Wheat | 1 | 2.69E-05 | 22 | 1 | 1 |
| QARQMVQAMRTIGTHP | InfluenzaA | 3 | 2.71E-19 | 22 | 4 | 0.75 |
| TTDPSFLGRY | SARS-CoV-2 | 18 | 2.27E-07 | 22 | 87 | 0.206896552 |
| YVYSRVKNL | SARS-CoV-2 | 2 | 1.99E-39 | 24 | 2 | 1 |
| NVILLNKHI | SARS-CoV-2 | 2 | 9.16E-13 | 24 | 2 | 1 |
| FRDYVDRFYKTLRAEQASQE | HIV-1 | 4 | 0.000569 | 24 | 27 | 0.148148148 |
| RLRAEAQVK | EBV | 18 | 1.35E-28 | 24 | 198 | 0.090909091 |
| NEGVKAAW | CMV | 18 | 1.37E-07 | 24 | 51 | 0.352941176 |
| MLDLQPETT | HPV | 18 | 1.26E-28 | 24 | 18 | 1 |
| GILGFVFTL | InfluenzaA | 19 | 0.001406 | 24 | 504 | 0.037698413 |
| SPRWYFYYL | SARS-CoV-2 | 20 | 6.85E-09 | 24 | 281 | 0.071174377 |
| NLVPMVATV | CMV | 23 | 1.65E-05 | 24 | 642 | 0.035825545 |
| KLGGALQAK | CMV | 30 | 0.000218 | 24 | 2190 | 0.01369863 |
| NCTFEYVSQPFLMDL | SARS-CoV-2 | 15 | 2.91E-23 | 29 | 81 | 0.185185185 |
| TFEYVSQPFLMDLE | SARS-CoV-2 | 18 | 1.60E-30 | 29 | 88 | 0.204545455 |
| TSQWLTNIF | SARS-CoV-2 | 2 | 1.99E-15 | 31 | 2 | 1 |
| YTVSCLPFT | SARS-CoV-2 | 3 | 1.63E-108 | 31 | 3 | 1 |
| YLQPRTFLL | SARS-CoV-2 | 42 | 3.24E-101 | 31 | 263 | 0.159695817 |
| KLGGALQAK | CMV | 65 | 0.001386 | 31 | 2190 | 0.029680365 |
| NCTFEYVSQPFLMDL | SARS-CoV-2 | 66 | 1.73E-24 | 31 | 81 | 0.814814815 |
| TFEYVSQPFLMDLE | SARS-CoV-2 | 69 | 1.16E-53 | 31 | 88 | 0.784090909 |
| RTLNAWVKV | HIV-1 | 2 | 5.70E-06 | 37 | 9 | 0.222222222 |
| MHQKRTAMFQDPQER | HPV-16 | 1 | 9.61E-24 | 42 | 1 | 1 |
| SPRWYFYYL | SARS-CoV-2 | 12 | 2.67E-13 | 42 | 281 | 0.042704626 |
| LLLEWLAMA | SARS-CoV-2 | 13 | 6.70E-08 | 42 | 13 | 1 |
| KLGGALQAK | CMV | 21 | 3.00E-18 | 42 | 2190 | 0.009589041 |
| NLSDRVVFV | SARS-CoV-2 | 1 | 0.00091 | 47 | 3 | 0.333333333 |
| NLSDRVVFV | SARS-CoV-2 | 1 | 0.00091 | 49 | 3 | 0.333333333 |
| KLGGALQAK | CMV | 10 | 0.00096 | 52 | 2190 | 0.00456621 |
| NQKLIANQF | SARS-CoV-2 | 9 | 2.52E-17 | 54 | 58 | 0.155172414 |
| KLGGALQAK | CMV | 18 | 1.69E-37 | 55 | 2190 | 0.008219178 |
| SLFNTVATLY | HIV-1 | 1 | 0.000455 | 60 | 1 | 1 |

| | | | | | | |
|---|---|---|---|---|---|---|
| KLGGALQAK | CMV | 9 | 0.001921 | 65 | 2190 | 0.004109589 |
| GPEPLPQGQLTAY | EBV | 1 | 0.000455 | 67 | 1 | 1 |
| LPEGLPQGQLTAY | EBV | 1 | 0.000455 | 67 | 1 | 1 |
| LPEPLGQGQLTAY | EBV | 1 | 0.000455 | 67 | 1 | 1 |
| LPEPLPQAQLTAY | EBV | 1 | 0.000455 | 67 | 1 | 1 |
| LPEPLPQGGLTAY | EBV | 1 | 0.000455 | 67 | 1 | 1 |
| LPEPLPQGQGTAY | EBV | 1 | 0.000455 | 67 | 1 | 1 |
| LPEPLPQGQLGAY | EBV | 1 | 0.000455 | 67 | 1 | 1 |
| LPEPLPQGQLTAY | EBV | 1 | 0.000455 | 67 | 1 | 1 |
| LPEPLPQGQLTGY | EBV | 1 | 0.000455 | 67 | 1 | 1 |
| RLQSLQTYV | SARS-CoV-2 | 1 | 0.000455 | 67 | 1 | 1 |
| TTDPSFLGRY | SARS-CoV-2 | 2 | 0.000406 | 78 | 87 | 0.022988506 |
| YLQPRTFLL | SARS-CoV-2 | 2 | 0.003623 | 85 | 263 | 0.007604563 |
| QLQPFPQPELPY | Wheat | 1 | 4.90E-15 | 87 | 1 | 1 |
| RAKFKQLL | EBV | 13 | 0.00048 | 87 | 239 | 0.054393305 |
| KLGGALQAK | CMV | 14 | 6.34E-07 | 87 | 2190 | 0.006392694 |
| NLVPMVATV | CMV | 4 | 0.002039 | 91 | 642 | 0.00623053 |
| RLNQLESKV | SARS-CoV-2 | 1 | 0.000455 | 99 | 1 | 1 |
| NLSDRVVFV | SARS-CoV-2 | 1 | 0.00091 | 108 | 3 | 0.333333333 |
| NLVPMVATV | CMV | 9 | 3.23E-08 | 110 | 642 | 0.014018692 |
| PTDNYITTY | SARS-CoV-2 | 9 | 5.45E-23 | 110 | 10 | 0.9 |

Supplementary table 2. Epitopes associated with TCRβ clusters for Cohort I.

| antigen.epitope | antigen.species | count | pval | cluster | all_clust_count | enrichment_score |
|---|---|---|---|---|---|---|
| TPQDLNTML | HIV-1 | 22 | 2.10E-05 | 0 | 25 | 0.88 |
| VTEHDTLLY | CMV | 33 | 5.01E-05 | 0 | 38 | 0.868421 |
| SIIAYTMSL | SARS-CoV-2 | 5 | 3.90E-05 | 3 | 5 | 1 |
| PKYVKQNTLKLAT | InfluenzaA | 17 | 2.86E-06 | 3 | 23 | 0.73913 |
| QARQMVQAMRTIGTHP | InfluenzaA | 18 | 0.000358 | 3 | 46 | 0.391304 |
| RLRAEAQVK | EBV | 19 | 3.63E-07 | 3 | 33 | 0.575758 |
| LTDEMIAQY | SARS-CoV-2 | 20 | 5.65E-06 | 3 | 24 | 0.833333 |
| NLVPMVATV | CMV | 78 | 2.86E-06 | 3 | 299 | 0.26087 |
| NEGVKAAW | CMV | 20 | 4.47E-06 | 4 | 21 | 0.952381 |
| TMADLVYAL | SARS-CoV-2 | 24 | 1.22E-10 | 4 | 24 | 1 |
| NLVPMVATV | CMV | 83 | 2.71E-06 | 4 | 299 | 0.277592 |
| RLRPGGKKK | HIV-1 | 3 | 0.001204 | 6 | 3 | 1 |
| VTEHDTLLY | CMV | 3 | 0.000347 | 7 | 38 | 0.078947 |
| LLYDANYFL | SARS-CoV-2 | 1 | 6.19E-05 | 11 | 3 | 0.333333 |
| RLNEVAKNL | SARS-CoV-2 | 3 | 6.51E-07 | 16 | 3 | 1 |
| SSGDATTAY | SARS-CoV-2 | 1 | 3.39E-05 | 54 | 1 | 1 |
| LLQTGIHVRVSQPSL | CMV | 4 | 2.21E-05 | 54 | 20 | 0.2 |

Supplementary table 3. Epitopes associated with both alpha and beta clusters for Cohort I. The odds and Pearson correlation scores for alpha-beta cluster co-occurrence is also shown.

| antigen.epitope | antigen.species | cluster_alpha | cluster_beta | odds | corr |
|---|---|---|---|---|---|
| RLRAEAQVK | EBV | 9 | 3 | 0.004513 | 0.49912 |
| RLRAEAQVK | EBV | 13 | 3 | 0.004578 | 0.361369 |
| RLRAEAQVK | EBV | 24 | 3 | 0.004357 | 0.585371 |
| RLRAEAQVK | EBV | 1 | 3 | 0.000902 | 0.338115 |
| NLVPMVATV | CMV | 9 | 3 | 0.004513 | 0.49912 |
| NLVPMVATV | CMV | 9 | 4 | 0.003094 | 0.473646 |
| NLVPMVATV | CMV | 24 | 3 | 0.004357 | 0.585371 |
| NLVPMVATV | CMV | 24 | 4 | 0.003049 | 0.550947 |
| NLVPMVATV | CMV | 110 | 3 | 0.004584 | 0.265179 |
| NLVPMVATV | CMV | 110 | 4 | 0.003117 | 0.280618 |
| NLVPMVATV | CMV | 0 | 3 | 0.001639 | 0.328225 |
| NLVPMVATV | CMV | 0 | 4 | 0.001598 | 0.314887 |
| NLVPMVATV | CMV | 91 | 3 | 0.004575 | 0.393508 |
| NLVPMVATV | CMV | 91 | 4 | 0.003117 | 0.360685 |
| SPRWYFYYL | SARS-CoV-2 | 9 | 3 | 0.004513 | 0.49912 |
| SPRWYFYYL | SARS-CoV-2 | 24 | 3 | 0.004357 | 0.585371 |
| SPRWYFYYL | SARS-CoV-2 | 0 | 3 | 0.001639 | 0.328225 |
| SPRWYFYYL | SARS-CoV-2 | 42 | 3 | 0.004567 | 0.552613 |
| NEGVKAAW | CMV | 24 | 4 | 0.003049 | 0.550947 |
| NEGVKAAW | CMV | 4 | 4 | 0.002123 | 0.572947 |
| TTDPSFLGRY | SARS-CoV-2 | 78 | 0 | 0.003347 | 0.115375 |
| TTDPSFLGRY | SARS-CoV-2 | 22 | 0 | 0.003316 | 0.529591 |
| TTDPSFLGRY | SARS-CoV-2 | 10 | 0 | 0.002674 | 0.620498 |
| TTDPSFLGRY | SARS-CoV-2 | 4 | 0 | 0.0022 | 0.613155 |
| LTDEMIAQY | SARS-CoV-2 | 0 | 3 | 0.001639 | 0.328225 |
| QARQMVQAMRTIGTHP | InfluenzaA | 22 | 3 | 0.00452 | 0.509571 |

Supplementary figure 1

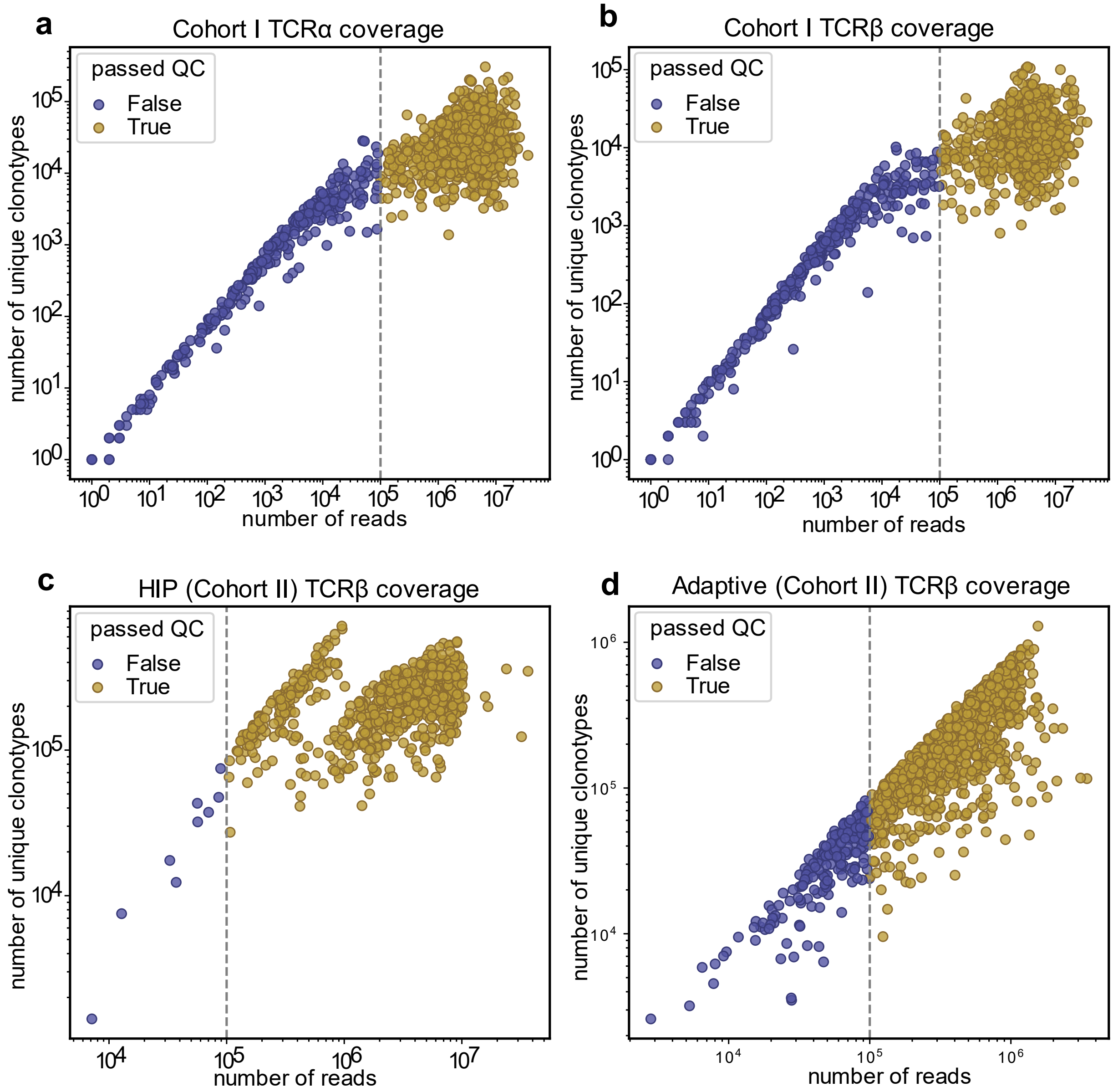


**Supplementary Figure 1. Distribution of sample coverage across technologies. A.** Sample coverage for Cohort I TCRα clonotypes. There were 1637 donors, 1273 samples left after preprocessing. **B.** The same as **A** but for TCRβ clonotypes. There were 1637 donors, 1225 samples left after preprocessing. For both β and α chains together, 1224 samples passed the read threshold. **C.** Distribution of coverage for TCRβ clonotypes in HIP cohort. There were 786 samples, but 9 were dropped during preprocessing. **D**. Distribution of coverage for TCRβ clonotypes in Adaptive cohort. There were 1414 samples, but 200 were dropped during preprocessing. HIP and Adaptive cohorts represent Cohort II.

Supplementary figure 2

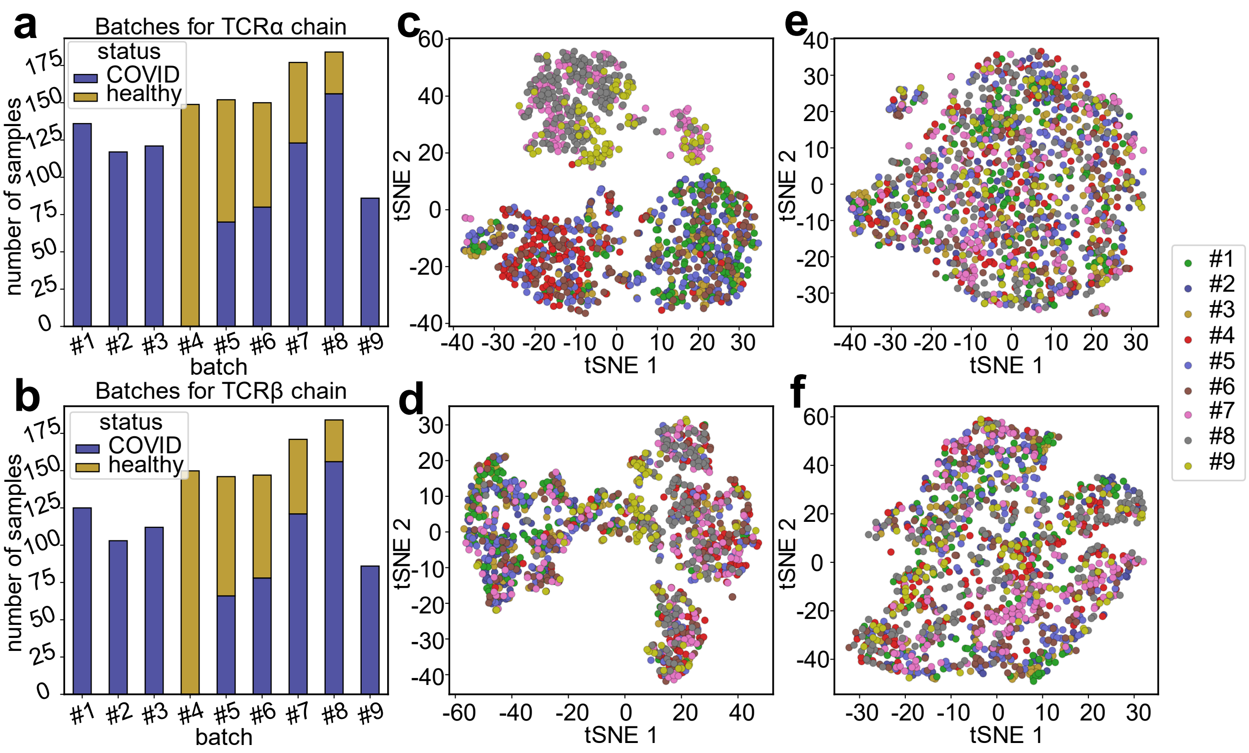


**Supplementary Figure 2. Additional information on data. A, B.** A bar plot of total number of subjects, number of COVID-19-convalescent and healthy cases for 9 batches of Cohort I reported in this study. The distribution is shown for samples with TCRα (**A**) and TCRβ (**B**) high-quality data. **C, D.** Visualization of batch effects using t-SNE. Plots show TRAV (**C**) and TRBV (**D**) gene usage profiles between samples before batch effect correction procedure, samples are colored by batch. **E, F**. TRAV (**E**) and TRBV (**F**) gene usage profiles after batch effect correction procedure, samples are colored by batch.

### Supplementary figure 3

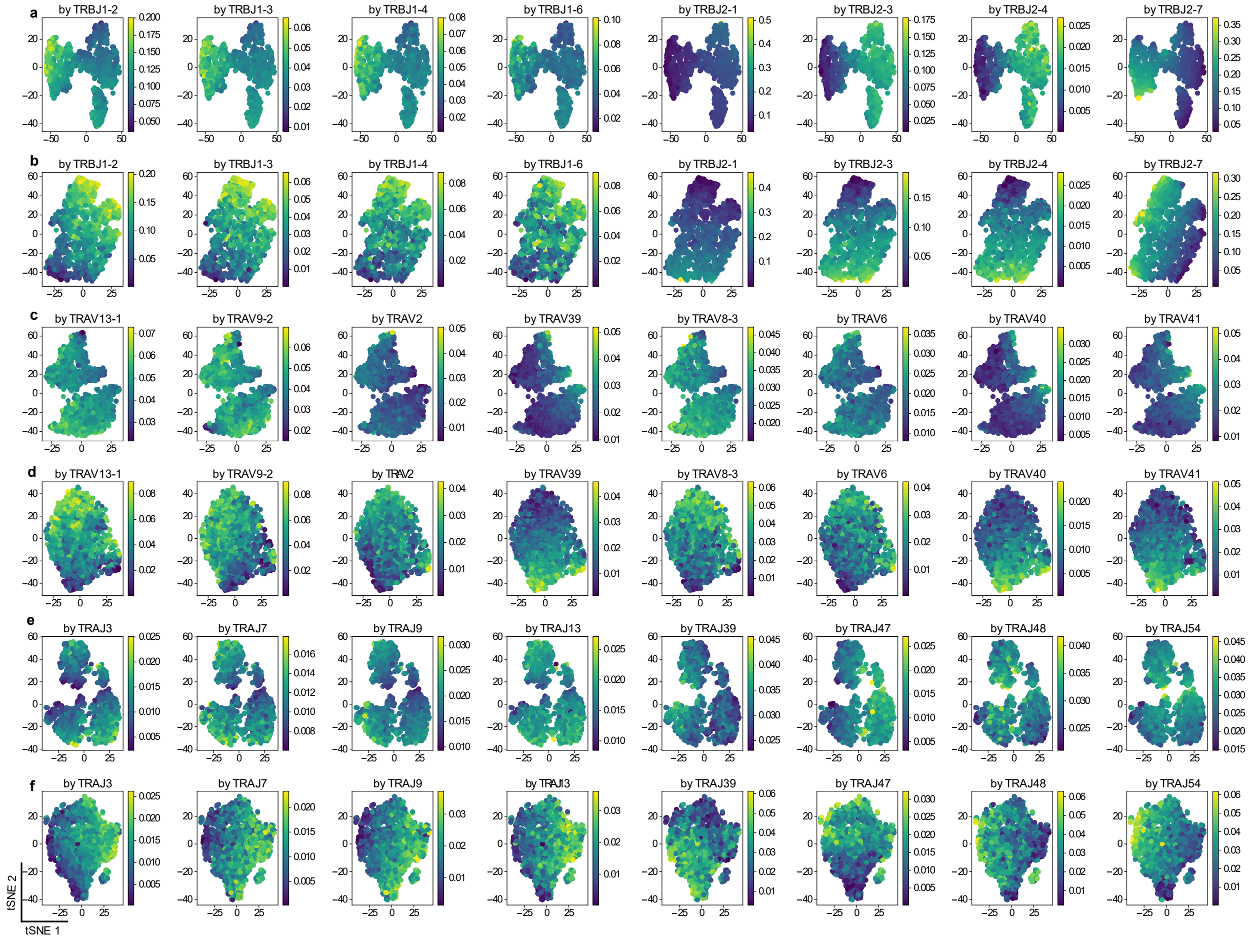


**Supplementary figure 3. Allele effects visualization for TCRβ J gene usages and TCRα V and J gene usages. A, B.** Visualization of biological effects pre (**A**) and post (**B**) batch effect correction procedure for TCRβ J gene usages. **C, D.** The same as **A, B**, but for TCRα V gene usages. **E, F.** The same as **A, B**, but for TCRα J gene usages.

### Supplementary figure 4

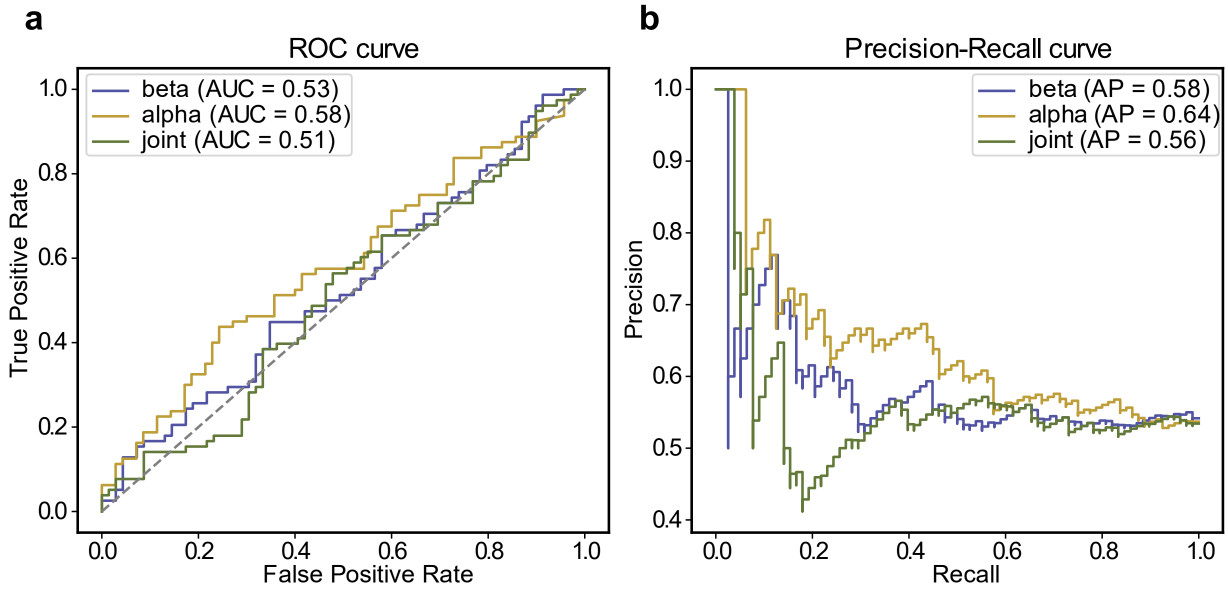


**Supplementary figure 4. COVID-19 status classification based on TCR α and β VJ gene usage. A.** ROC Curve showing three classifier scores. α corresponds to the classifier based on TRAV and TRAJ usages. β corresponds to TRBV and TRBJ usages. Joint is the classifier which uses both TCR α and β genes. **B.** The same as A, but for precision-recall curve.

### Supplementary figure 5

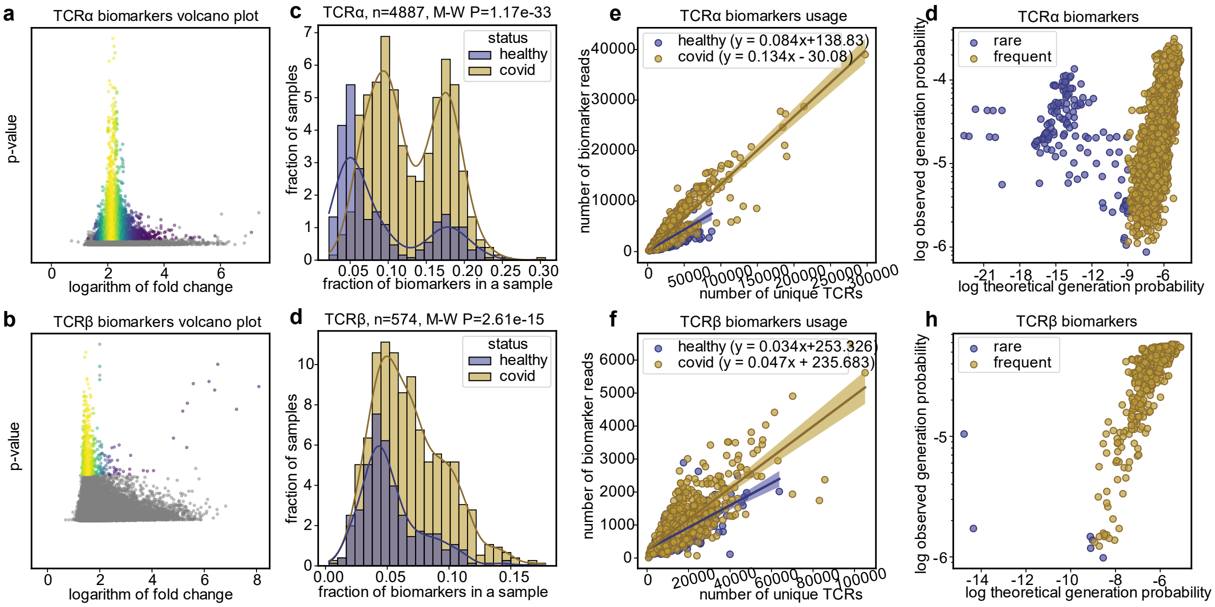


**Supplementary Figure 5. Analysis of COVID-19 TCR α and β biomarkers before cleanup procedure. A, B.** Volcano plot representing the position of selected clonotypes among all the clonotypes of interest for TCRα

(**A**) and TCRβ (**B**) chain. **C, D.** Scatter plot showing the frequency of COVID-19-associated TCRα (**C**) and TCRβ (**D**) clonotypes plotted against the number of unique TCRβ clonotypes in Cohort I before the cleanup. **E, F.** Distribution of COVID-associated TCRα (**E**) and TCRβ (**F**) clonotypes across healthy and convalescent samples before the cleanup. **G, H.** Scatterplot showing the usage of TCRα (**G**) and TCRβ (**H**)clonotypes in samples against the expected frequency (calculated using probability based model software OLGA).

#### Supplementary figure 6

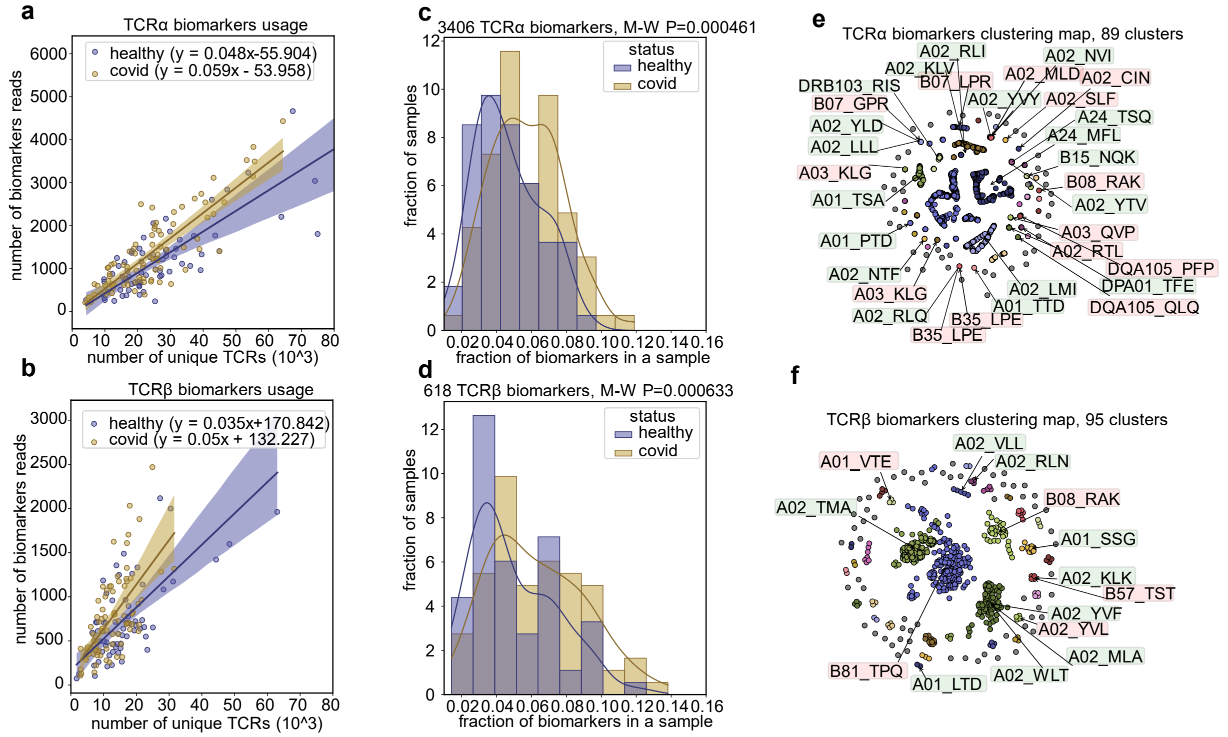


**Supplementary Figure 6. Biomarkers analysis for statistical testing without a test batch (#6). A, B.** Scatter plot showing the frequency of COVID-19-associated TCRα (**A**) and TCRβ (**B**) clonotypes plotted against the number of unique clonotypes in the Cohort I for the dropped batch (#6). **C, D.** Distribution of COVID-associated TCRα (**C**) and TCRβ (**D**) clonotypes across healthy and convalescent samples for the dropped batch (#6). **E, F.** TCRα (**E**) and TCRβ (**F**) clonotype clustering. The epitopes marked with green are the SARS-CoV-2 epitopes, while the red ones are the epitopes belonging to other viruses.

#### Supplementary figure 7

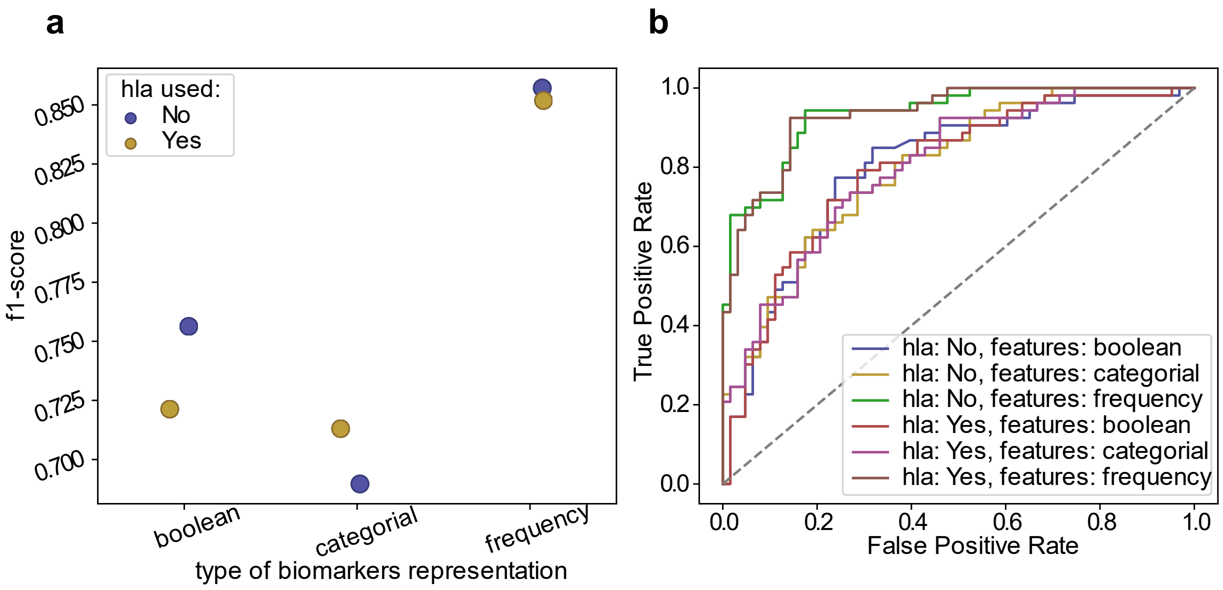


**Supplementary Figure 7. Comparison of model settings for joint meta-classifiers. A.** F1 score of models trained on boolean/categorial/frequency features. The frequency features are gaining the highest f1-scores. **B.** Roc-curve, showing that frequency feature-based classifiers are getting the highest scores, whereas the HLA boolean features do not contribute to the model performance significantly.

#### Supplementary figure 8

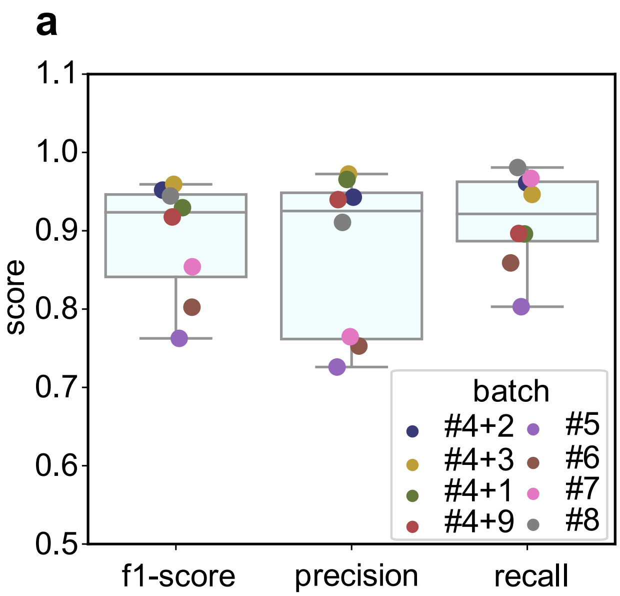


**Supplementary figure 8. TCRα+β metaclonotype model cross validation between batches. A.** Evaluation of target metrics (f1-score, precision, recall) for one batch out cross validation. Each point represents the target metrics for one of the batches. Each batch is thrown out as a validation set and the entire model is being trained using the rest of the data. Joint metaclonotype model is used here. The error bars represent the range of metrics between thrown out batches. One can see that the summarized batches result in a better quality.

#### Supplementary figure 9

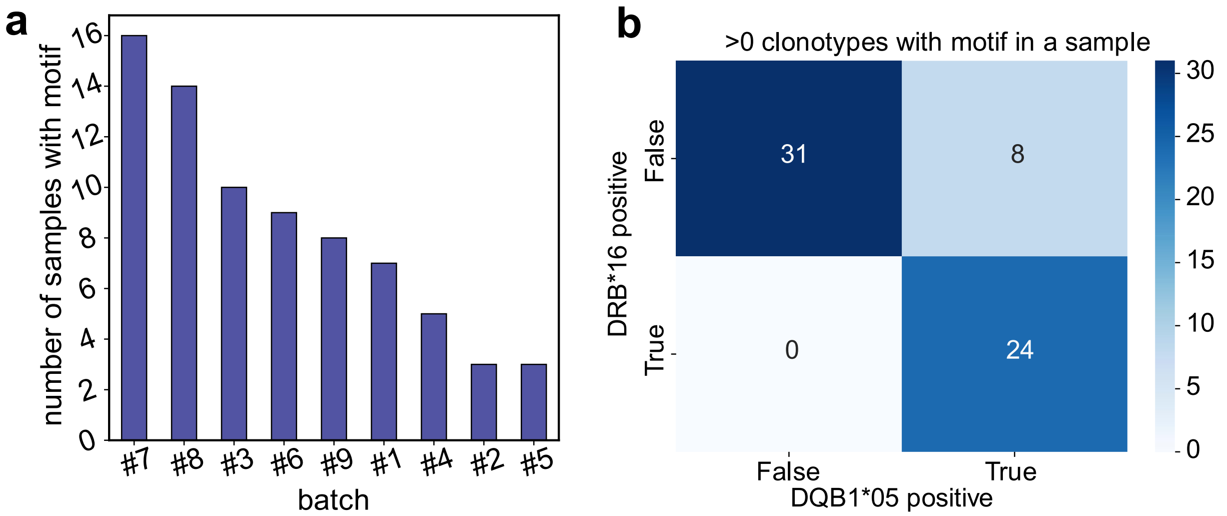


**Supplementary Figure 9**. **Heatmaps of HLA-DRB1*16 and HLA-DQB1*05 presence in COVID-19 patients with clonotypes forming the CASSRTGXGSSYNSPLHF pattern. A.** Distribution of samples with motif between batches. The motif clonotypes are present in all the batches which demonstrates the absence of batch effect. **B.** Heatmap representing the linkage of HLAs of interest for all the COVID-19 patients with at least one read corresponding to pattern clonotypes of interest.

#### Supplementary figure 10

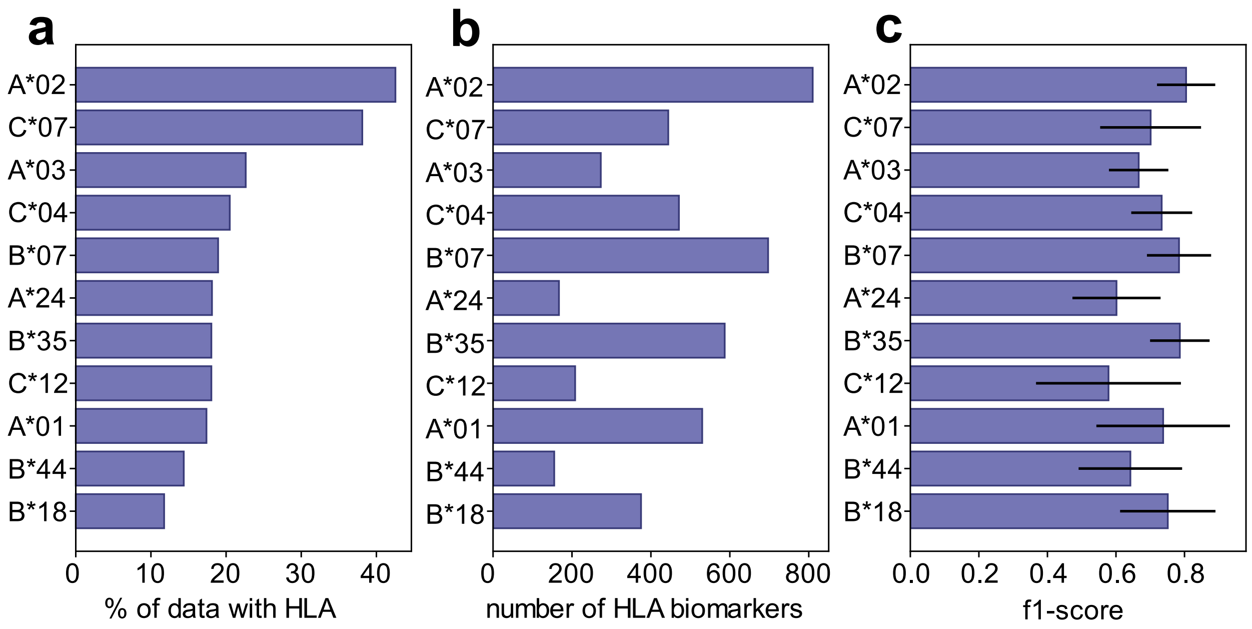


**Supplementary figure 10. Predicting donor HLA allele based on HLA-associated clonotypes. A.** Number of HLA positive donors in Cohort I. **B.** Number of clonotypes associated with presence of HLA for each allele.

**C.** Average F1-scores for one-batch-left-out cross validation. Error bars show 25-75% quartiles for F1 score of each thrown out batch.

#### Supplementary figure 11

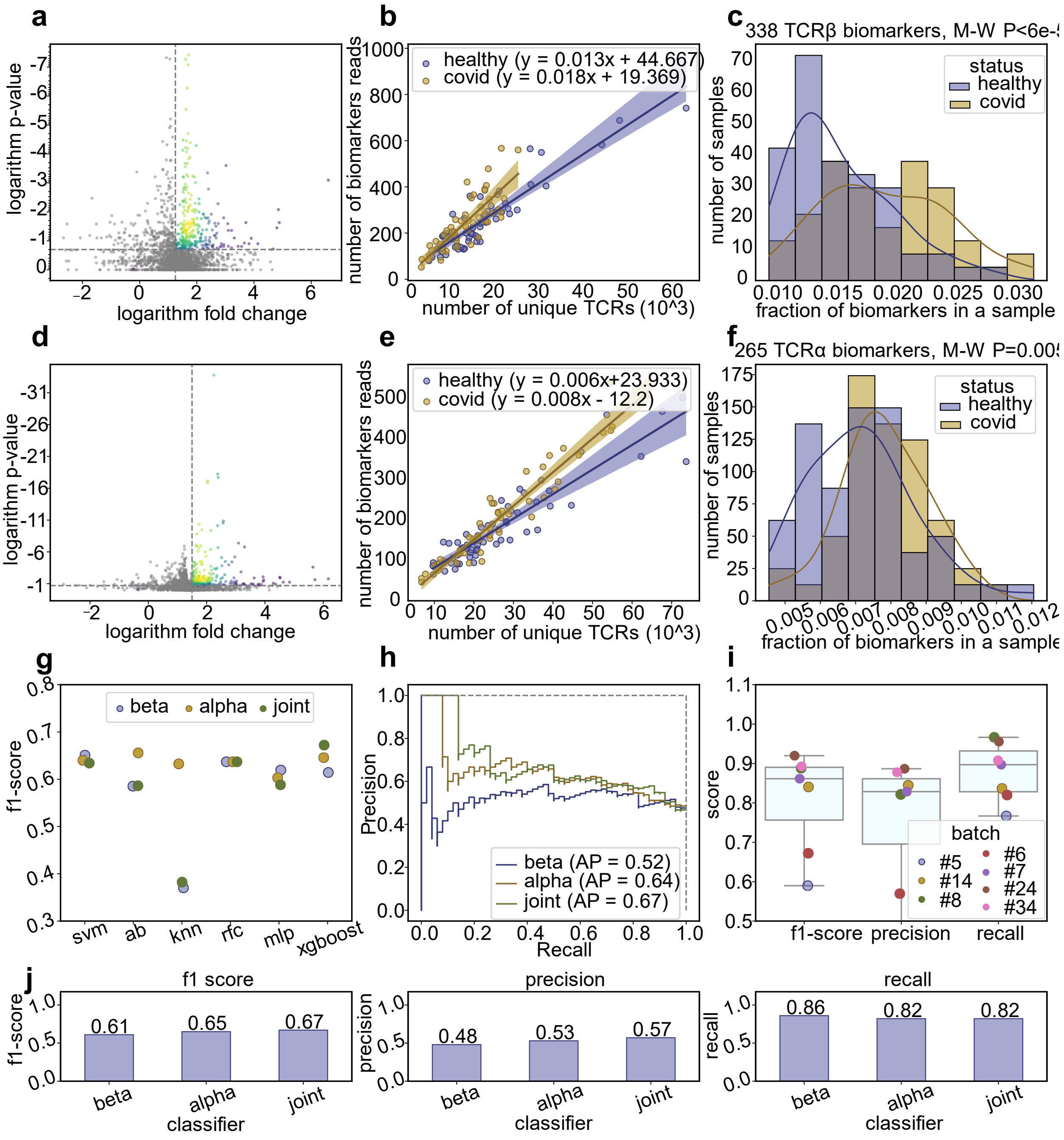


**Supplementary Figure 11. Analysis of models built using VDJDb TCR α and β SARS-CoV-2 clonotypes.** **A.** Volcano plot for VDJDb TCRβ SARS-CoV-2 clonotypes. The fold change of clonotype usage between healthy and convalescent cohorts in comparison to Fisher exact test p-value is shown. The dashed lines show the selected thresholds. Multiple hypothesis correction was not applied. **B.** Scatter plot showing the frequency of VDJDb TCRβ clonotypes plotted against the number of unique TCRβ clonotypes. **C.** Distribution of

COVID-associated TCRβ clonotypes across healthy and convalescent samples. **D, E, F.** The same as **A, B, C,** but for VDJDb TCRα clonotypes. **G.** Distribution of f1-scores for biomarker sets across different models. SVM shows most stable results. **H.** The precision-recall curve showing the difference in model performance. **I.** Evaluation of target metrics (f1-score, precision, recall) for one-batch-left-out cross validation. The metrics were evaluated for the TCR α+β biomarkers set. The boxplots represent the distribution of scores between different evaluation batches. **J.** Evaluation of target metrics for different biomarker sets.
